# *Candida albicans* enhances melanoma cell aggressiveness through p38-MAPK and HIF-1α pathways and metabolic reprogramming

**DOI:** 10.1101/2025.01.17.633543

**Authors:** Leire Aparicio-Fernandez, Nahia Cazalis-Bereicua, Maialen Areitio, Oier Rodriguez-Erenaga, Lucia Abio-Dorronsoro, Leire Martin-Souto, Idoia Buldain, Joana Márquez, Aitor Benedicto, Beatriz Arteta, Nuria Macias-Cámara, Monika Gonzalez, Jose Ezequiel Martin Rodriguez, Ana Aransay, Aize Pellon, David L. Moyes, Juan Anguita, Aitor Rementeria, Aitziber Antoran, Andoni Ramirez-Garcia

## Abstract

Recent studies have increasingly focused on the role of fungi, including *Candida albicans*, in carcinogenesis. Since *C. albicans* is a component of the human microbiota, particularly on the skin, we investigated its effect on the phenotype and signalling pathways of melanoma cells. Assays for migration, adhesion, angiogenesis, and hepatic metastasis showed that *C. albicans* promotes a more malignant phenotype in melanoma cells. At the transcriptomic level, *C. albicans* increased the expression of VEGF (*Vegfa*), and genes associated with MAPK and HIF-1 signalling pathways, and with aerobic glycolysis. Further in vitro analysis revealed that TLRs and EphA2 receptors are involved in the recognition of live *C. albicans*, stimulating VEGF secretion and expression of the AP-1 transcription factor component c-Fos through p38-MAPK and HIF-1α. These pathways also regulate the expression of other AP-1 constituents such as *Atf3*, *Jun*, and *Jund*. Moreover, p38-MAPK regulates glycolytic genes like *Hk2*, *Slc2a1*, and *Eno2*. In conclusion, *C. albicans* activates the p38-MAPK/c-Fos/AP-1 and HIF-1/HIF-1α/c-Fos/AP-1 pathways in melanoma cells, promoting a pro-angiogenic environment and metabolic reprogramming. Therefore, this study clarifies the impact of *C. albicans* on melanoma cells, which can lead to the use of antifungal therapies as complementary to traditional treatments for melanoma.

## 1. Introduction

Cancer is one of the main causes of death worldwide, resulting in approximately 10 million deaths in 2020 alone ^1^. It is estimated that between 13% and 18% of these cases are caused by microorganism infections ^2,3^. In fact, the International Agency for Research on Cancer (IARC) has already classified thirteen microorganisms as carcinogenic, including viruses, bacteria, and parasites ^4^. However, recent studies have shown that other microorganisms could also directly promote cancer. While most studies focus on virus and bacteria interactions with cancer, some research also highlights a link between cancer progression and certain fungi, such as *Malassezia*, *Aspergillus fumigatus* and *Candida albicans* ^5–8^.

*C. albicans* is an opportunistic dimorphic fungus that is usually part of the human microbiota in many locations, such as the oral cavity, gastrointestinal tract, or vagina ^9,10^. However, in the last decades, it has been suggested that it could promote cancer initiation and progression. Cawson and Williamson in 1969 were the first who related *Candida* presence with promotion of cancer, specifically leukoplakia, which is a precancerous oral lesion ^11,12^. Since then, many studies have described its association especially with oral and oesophageal cancer in humans ^13–16^, but also with other types of cancers and metastasis ^6–8,17^. Nevertheless, the mechanisms by which *C. albicans* can induce cancer progression and metastasis have not been completely elucidated. It has been proposed that production of carcinogenic compounds such as acetaldehyde or nitrosamine, chronic inflammation, or induction of metastasis could play a role ^18,19^.

In this context, it has been demonstrated that *C. albicans* can induce cellular processes associated with cancer including enhanced cell migration, the production of oncometabolites, and the activation of pro-inflammatory cytokines and the expression of cancer-related genes in various cell lines *in vitro* ^8,16,20–22^. Moreover, *in vivo* oral cancer models have also linked *C. albicans* to augmented cancer development and progression ^16,23,24^. These results are supported by *in vivo* research conducted by our group employing melanoma cells, a particularly aggressive cancer type, which revealed that *C. albicans* increases liver metastasis and creates a pro-metastatic microenvironment characterised by an enhanced release of Tumour Necrosis Factor (TNF) and Interleukin-18 (IL-18) ^25–27^. The stimulation of pro-tumour and pro-metastatic pathways in melanoma and liver cells by this fungus may be the direct cause of this pro-metastatic action. However, despite the potential for *C. albicans* and melanoma cells to interact within the skin or bloodstream, where both can disseminate and coexist, the specific impact of *C. albicans* on melanoma cell aggressiveness has yet to be explored.

As *C. albicans* is a microorganism frequently isolated from the skin, we aimed to examine the role that this opportunistic fungus plays in promoting the development of cancer in melanoma cells. Using *in vitro* and *in vivo* techniques we demonstrate that *C. albicans* induces phenotypic changes and metabolic reprogramming in melanoma cells, increasing their aggressiveness through activation of inflammatory, pro-angiogenic, and hypoxic pathways, specifically p38-MAPK/c-Fos and HIF-1α/c-Fos. These findings support the development of new therapies for melanoma cases influenced by *C. albicans*.

## 2. Results

### 2.1. C. *albicans stimulates migration, adhesion, pro-angiogenesis, and hepatic metastasis, but no proliferation, in melanoma cells*

*In vitro* assays revealed that heat-killed (HK)-*C. albicans* yeasts promoted a significant increase in the migration ability of melanoma cells (**Fig. 1a**) and their adhesion capacity to liver sinusoidal endothelial cells (LSECs) (**Fig. 1b**). Furthermore, a significant increase in the migration capacity of LSECs was observed when they were incubated with conditioned media obtained from melanoma cells exposed to live *C. albicans*, which could indicate the presence of pro-angiogenic substances in the media (**Fig. 1c**). However, HK-*C. albicans* yeast failed to increase melanoma cell proliferation (**Suppl. Fig. 1**).

**Figure 1.**
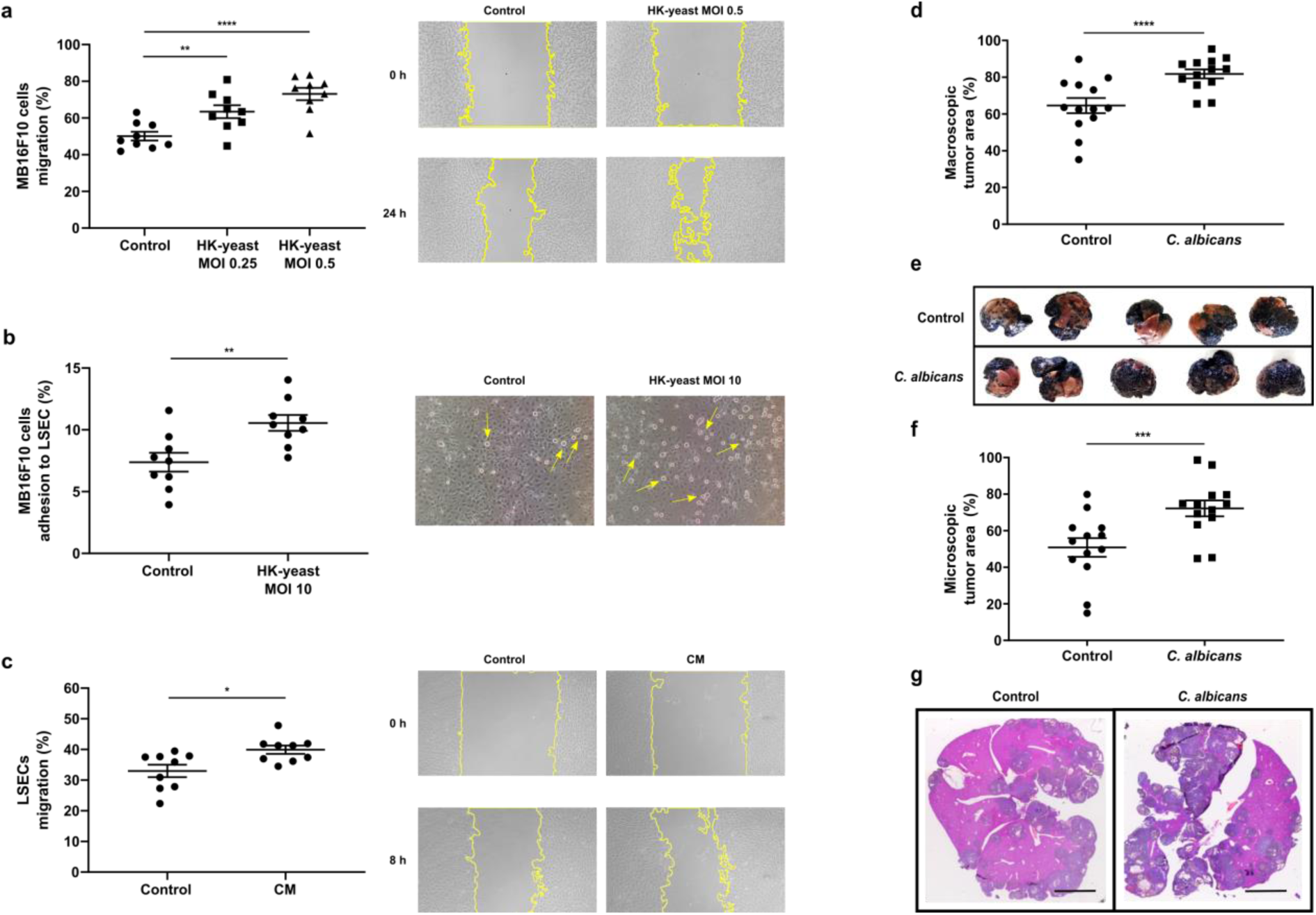
*Candida albicans* effect in melanoma cells phenotype. **a** Melanoma cells migration after 24 hours of exposure to HK-*C. albicans* yeast MOI of 0.25 and 0.5. **b** Melanoma cells adhesion to LSECs after 24 hours of exposure of melanoma cells to HK-*C. albicans* yeast MOI 10. **c** LSECs migration after eight hours of exposure to the conditioned media collected from melanoma cells exposed to live *C. albicans* MOI 5 of the fungus for six hours. **a**, **b** and **c** graphs represent all the results obtained and images (10X magnification) are representative of one experiment of each assay (*n* = 9 technical replicates from 3 biologically independent samples). Effect of *C. albicans* stimulation of melanoma cells in liver metastasis promotion (*n* = 13 biologically independent samples): **d** Liver area occupied by metastatic melanoma in the liver by control group and *C. albicans*-stimulated melanoma cells and **e** representative macroscopic liver images showing an augmented size of metastatic area in six hours *C. albicans*-stimulated melanoma cells injected group. **f** Percentage of the microscopic tumour area in the liver originated by control group and *C. albicans*-stimulated melanoma cells and **g** representative H&E staining showing an increased metastatic colonization of melanoma in *C. albicans*-stimulated melanoma cells injected group. Scale bar 500 μm. For all data, individual values and mean ± SEM are shown. *, **, and *** denote *P* < 0.05, *P* < 0.01, and *P* < 0.001, respectively (two-tailed, unpaired, *t-student* test). HK: Heat-Killed, CM: Conditioned Media.

In addition to the *in vitro* assays, the pro-metastatic effect induced by *C. albicans* was determined by intrasplenic inoculation in mice of melanoma cells previously stimulated with live *C. albicans*. After 14 days, all mice injected with melanoma cells developed hepatic metastasis (**Fig. 1d, f**), but both macroscopic (**Fig. 1d, e**) and microscopic (**Fig. 1f, g**) tumour areas were augmented in the livers of mice inoculated with *C. albicans*-stimulated melanoma cells compared to those inoculated with non-stimulated cells.

### 2.2. *Transcriptional response of melanoma cells to* C. *albicans stimulation*

The study of the mechanisms underlying the alterations of the phenotype induced by *C. albicans* was first carried out through a transcriptomic analysis of melanoma cells exposed to live fungal cells (Multiplicity of infection (MOI) 1) for six hours. The results showed 63 genes significantly (│FC│ > 1.5, p_adj_ < 0.05) upregulated while no genes were downregulated in comparison to melanoma cells not exposed to the fungus (**Fig. 2a, Suppl. Table 1**). Notably, among the differentially-expressed genes (DEGs), genes involved in signalling pathways (*Dusp1*, *Dusp8*, *Trib3*), glucose metabolism (*Hk2*, *Slc2a1*, *Pdk1*), and transcription factors (*Jun*, *Fos, Atf3*) were upregulated. In addition, 54% of the genes (34 out of 63) and nine of the first 15 Gene Ontology-Biological Processes (GO-BPs; ordered according to the GeneRatio) were associated with a response to stress conditions, of which four GO-BPs were specifically related to hypoxia (**Fig. 2b**). With respect to KEGG analysis, only three pathways were significantly upregulated in response to *C. albicans*: MAPK signalling pathway (mmu04010), HIF-1 signalling pathway (mmu04066), and renal cell carcinoma (mmu05211) (**Fig. 2c**). Interestingly, these three pathways share the gene *Vegfa,* which codes for the pro-angiogenic factor VEGF (**Fig. 2d**).

**Figure 2.**
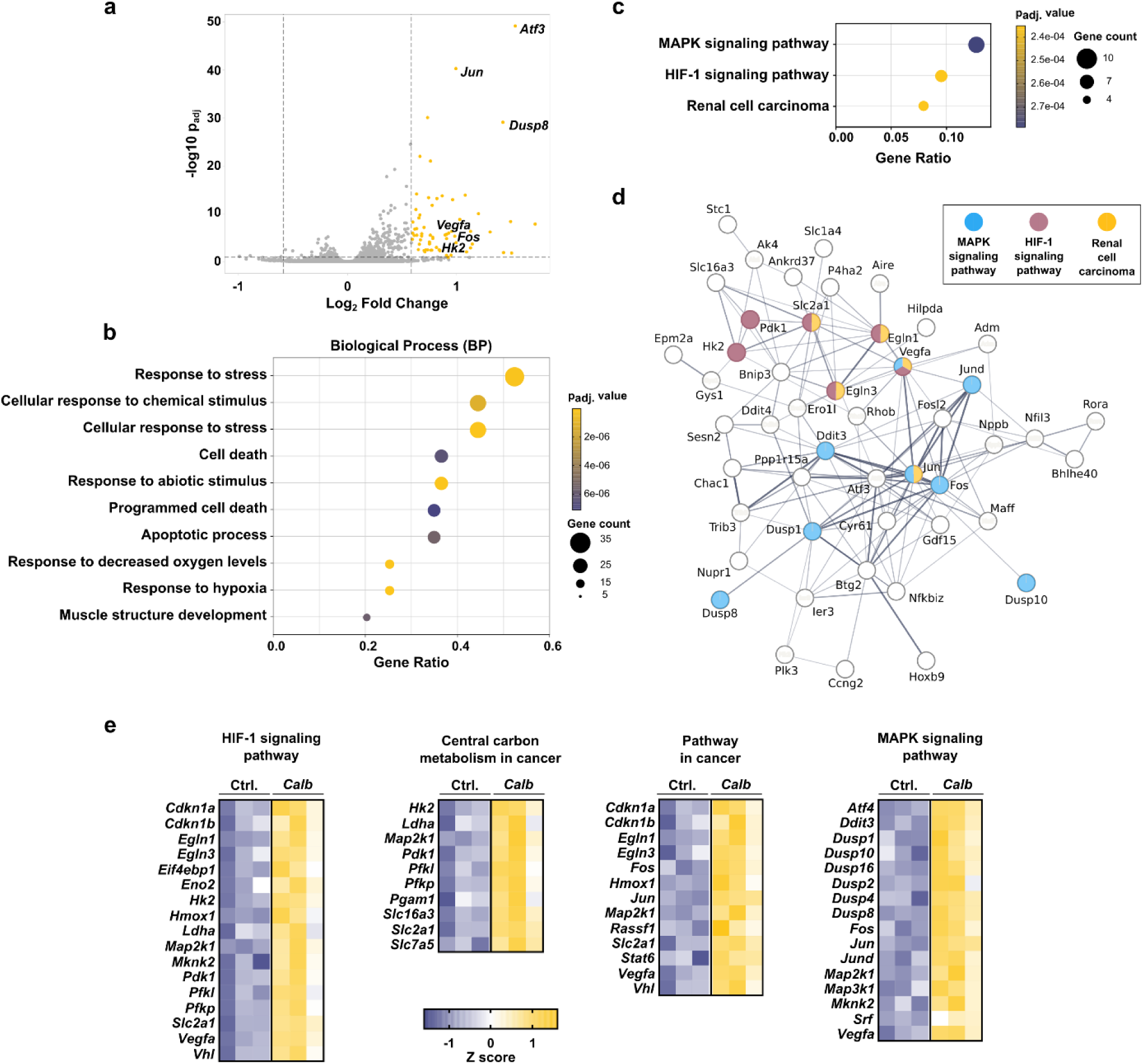
Transcriptomic analysis of melanoma cells exposed to live *Candida albicans* at MOI 1 for six hours. **a** Volcano plot representing DEGs. Yellow dots represent upregulated genes. **b** GO enrichment analysis of the Biological Processes (GO-BP) significantly upregulated. **c** KEGG pathways significantly upregulated. **d** STRING performed with upregulated genes, their relationship (grey lines) and the KEGG pathways in which they are implicated (only interrelated genes are shown). Line thickness indicates the strength of data support. **e** Heat-maps showing the expression levels comparison between samples (Z-score) of DEGs: HIF-1 signalling pathways, central carbon metabolism in cancer, pathways in cancer and MAPK signalling pathways. Each column represent a biologically independent sample. Yellow and purple boxes represents upregulated and downregulated genes, respectively. **a**, **b**, **c** and **d** analysis were performed for │FC│ > 1.5 and p_adj_ ≤ 0.05 and, **e** analysis with p_adj_ ≤ 0.05 (iDEP 1.0 tool and Deseq2 method) (*n* = 3 biologically independent samples). DEGs: differentially-expressed genes.

A second less restrictive bioinformatics analysis of the results was performed, this time only considering the p_adj_ (< 0.05) as a selecting parameter to give a more extensive view of the pathways activated. Using this parameter, 271 upregulated genes (**Suppl. Table 2**) were detected, several of them being involved in the HIF-1 signalling pathways, central carbon metabolism in cancer (mmu05230), pathways in cancer (mmu05200), and MAPK signalling pathways. Of note, gene expression of some kinases (*Map2k1*, *Map3k1* or *Mknk2*), signalling regulators (e.g. *Dusp2*, *Dusp4*) or glycolytic enzymes (e.g. *Eno2*, *Ldha*) were found to be upregulated using these settings (**Fig. 2e**).

### 2.3. C. *albicans drives a response in melanoma cells that leads to VEGF release*

We next investigated the effects of fungal morphology, viability, and conditioned media on melanoma cells in greater detail. For that, the release of the pro-angiogenic molecule VEGF and the expression of the *Fos* gene, an important MAPK-related transcription factor overexpressed in the transcriptomic analysis, were quantified by ELISA and RT-qPCR respectively (**Fig. 3a, b**). The highest levels of VEGF release (**Fig. 3a**) and *Fos* expression (**Fig. 3b**) were observed after the co-incubation of cancer cells with live *C. albicans*. The rest of the conditions promoted a lesser, statistically non-significant increase in VEGF, and barely altered the expression of the *Fos* gene. Thus, for subsequent experiments only live *C. albicans* was used.

**Figure 3.**
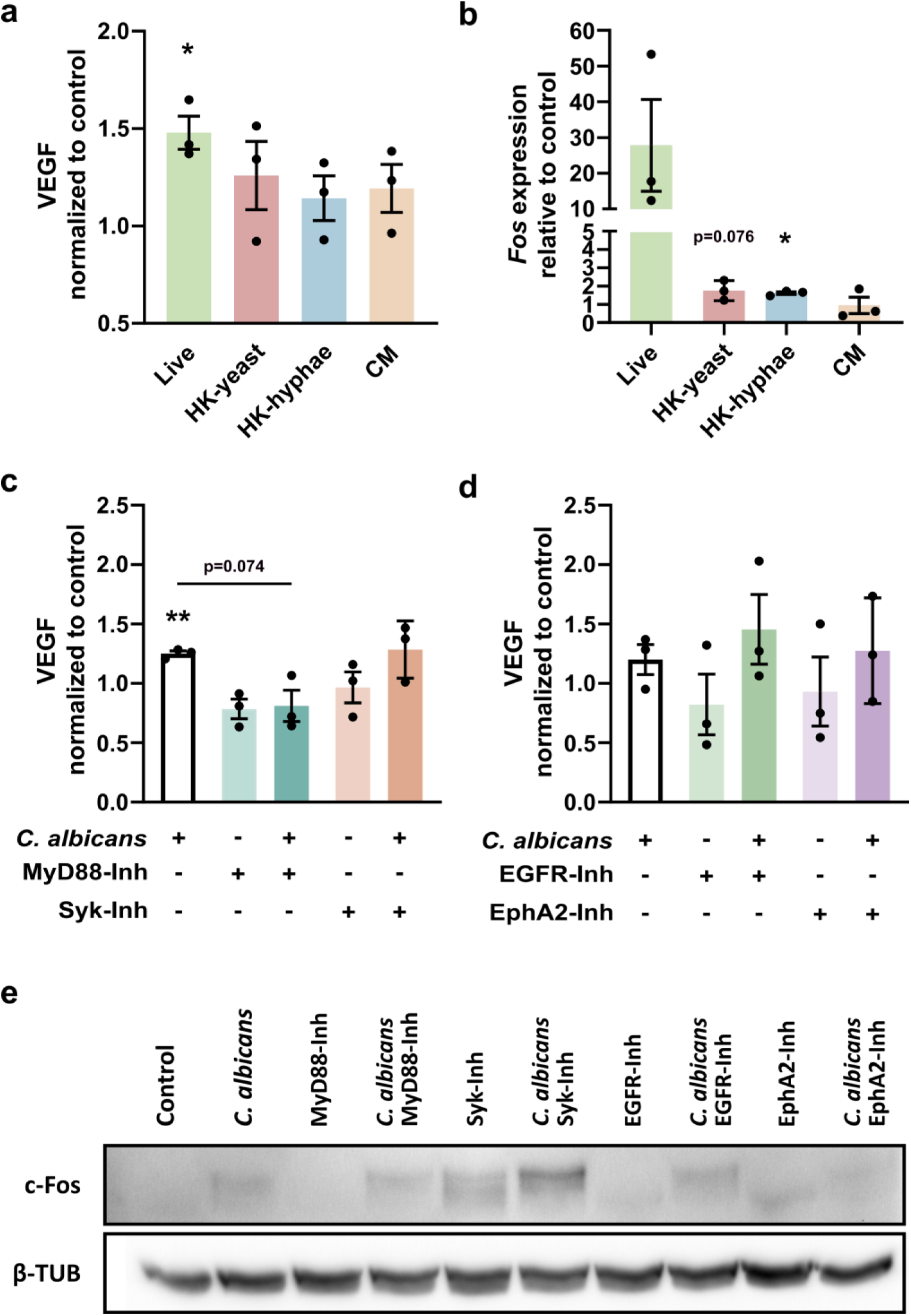
Influence of the viability of *Candida albicans* and the main pathogen recognition receptors (PRRs) in melanoma cells after six hours of stimulation. a VEGF cytokine release and b *Fos* gene expression induced by different morphologies and released molecules of *C. albicans*. **c**, **d** VEGF cytokine release and **e** c-Fos protein expression in live *C. albicans*-exposed melanoma cells alone or treated with inhibitors for **c**, **e** TLRs (MyD88 inhibitor: TJ-M2010-5, 30 µM) and CLRs (Syk inhibitor: Piceatannol, 30 µM), and **d**, **e** EGFR (PD153035, 1 µM) and EphA2 (NVP-BHG712, 1 µM). **a**, **b**, **c**, **d** Individual values and mean ± SEM are shown (*n* = 3 biologically independent samples). * and ** denote *P* < 0.05 and *P* < 0.01 in comparison to control without *C. albicans*, respectively (two-tailed, unpaired, *t-student* test). **e** Representative data of three independent Western Blots. HK: Heat-Killed; CM: Conditioned Media obtained after six hours of exposure of melanoma cells to live *C. albicans* MOI 5.

To investigate the initial host-fungus contact that activates the melanoma cell response, the role of the primary receptors involved in host-*Candida* interactions, mainly pattern recognition receptors (PRRs), were studied. Molecular inhibitors for the receptors EGFR and EphA2, and for the signal transducing proteins for almost all TLRs (MyD88) and CLRs (Syk), were used to assess their effects on VEGF release and c-Fos protein expression (**Fig. 3c-e**). The inhibition of MyD88-mediated TLRs signal transduction resulted in a significant reduction of VEGF release induced by *C. albicans* but did not show a significant effect on c-Fos protein expression, which appeared to be mainly mediated by the β-glucan receptor EphA2. However, inhibition of both Syk and EGFR did not alter any of the two parameters analysed.

### 2.4. *The release of VEGF by melanoma cells in response to* C. *albicans is mediated by p38-MAPK and HIF-1*α

To decipher the signal cascade induced by *C. albicans* in melanoma cells, the impact on VEGF release of the three main pathways activated by MAPK (ERK1/2, JNK and p38), and the inflammation-related NF-κB pathway, was studied using different inhibitors for p38, JNK, ERK1/2 and NF-kB (**Fig. 4a**). The results demonstrated a significant reduction in VEGF release in cells stimulated by *C. albicans* when p38-MAPK was inhibited, whereas no alterations were observed with the inhibitors for ERK1/2, JNK, or NF-κB.

**Figure 4.**
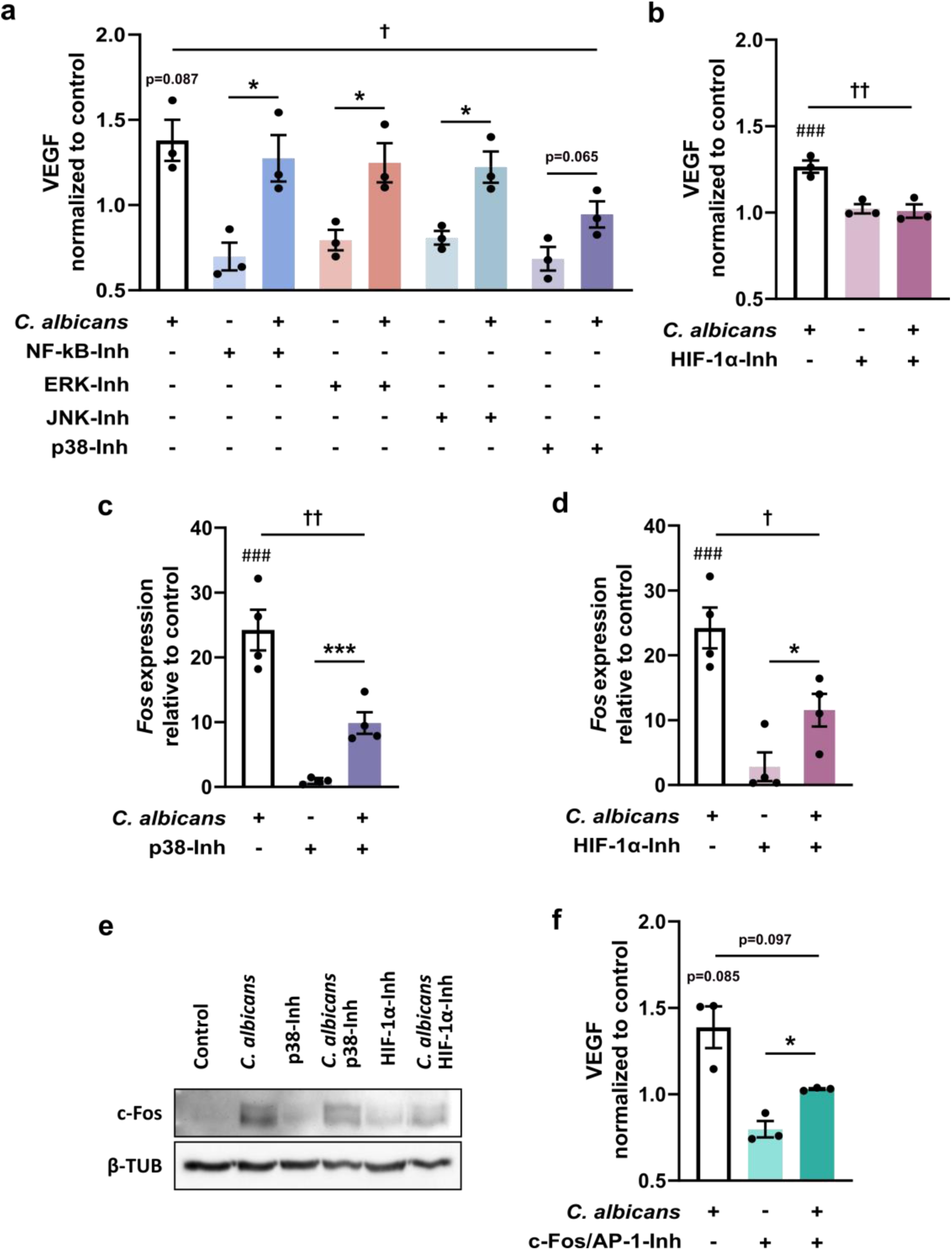
Live *Candida albicans* at MOI 5 regulate MAPK and HIF-1α pathways in melanoma cells. VEGF cytokine release induced by six hours of stimulation with *C. albicans* alone or treated with inhibitors for **a** NF-κB (BAY 11-7082, 2 µM), ERK1/2 (PD-0325901; 1 µM), JNK (SP600125, 10 µM), p38 (SB 203580, 10 µM), **b** HIF-1α (LW6, 10 μM) and **c** c-Fos/AP-1 complex (T-5224, 10 µM). **c**, **d** *Fos* gene and **e** protein expression analysis alone or treated with p38 inhibitor or HIF-1α inhibitor. **f** VEGF cytokine release induced by six hours of stimulation with *C. albicans* alone or treated with inhibitors for c-Fos/AP-1 complex (T-5224, 10 µM). Individual values and mean ± SEM are shown. **a**, **b** and **f** (*n* = 3 biologically independent samples); **c** and **d** (*n* = 4 biologically independent samples). ^###^ denotes *P* < 0.005 respect to control without *C. albicans*; * and *** denote *P* < 0.05 and *P* < 0.005 of each inhibitor compared to inhibitor with *C. albicans*, respectively; †, †† and ††† denote *P* < 0.05, *P* < 0.01 and *P* < 0.005 of *C. albicans* group compared to inhibitors with *C. albicans* groups, respectively (two-tailed, unpaired, *t-student* test). **e** Representative data of three independent Western Blots. HK: Heat-Killed; CM: Conditioned Conditioned Media obtained after six hours of exposure of melanoma cells to live *C. albicans* MOI 5.

The involvement of the HIF-1 signalling pathway, identified as the second pathway in the KEGG analysis, in VEFG release was then studied using a specific HIF-1α inhibitor (**Fig. 4b**). The data revealed a significant reduction in VEGF release in cells stimulated with *C. albicans* when treated with the inhibitor, compared to those without inhibitor treatment, indicating that, in addition to p38-MAPK, HIF-1α regulates VEGF release induced by *C. albicans*.

Previous work has identified c-Fos as a central component in the MAPK signalling circuit activated by *C. albicans* infection of epithelial cells ^28,29^. Thus, we next set out to determine whether p38-MAPK and HIF-1α inhibition impacted on c-Fos gene and protein expression in melanoma cells. Inhibition of both pathways demonstrated a significant reduction of *Fos* gene and c-Fos protein expression in cells infected with *C. albicans* (**Fig. 4c-e**). However, neither of them alone reduced its gene and/or protein expression to control levels, suggesting that they are independently involved in c-Fos production.

Given the central role played by p38-MAPK in regulating VEGF release, we investigated whether activation of the MAPK-activated c-Fos/AP-1 transcription factor component plays a role in VEGF production and release (**Fig. 4f**). As expected, the inhibition of the c-Fos-containing AP-1 transcription factor (c-Fos/AP-1) resulted in a decrease in the VEGF release in the presence of *C. albicans* with respect to the cells stimulated with the fungus without the inhibitor, confirming that the c-Fos/AP-1 complex is, at least in part, responsible for VEGF production.

### 2.5. *The release of VEGF by melanoma cells in response to* C. *albicans is mediated by the AP-1 transcription factor*

The dimeric transcription factor AP-1 is formed by proteins from the Fos, Jun, ATF, and MAF families ^30^. Our transcriptomic analysis revealed that, in addition to *Fos*, *C. albicans* exposure induced the expression of the *Atf3*, *Jund*, and *Jun* genes. Thus, the involvement of the p38-MAPK and HIF-1α in regulating the activation of these genes was investigated (**Fig. 5**).

**Figure 5.**
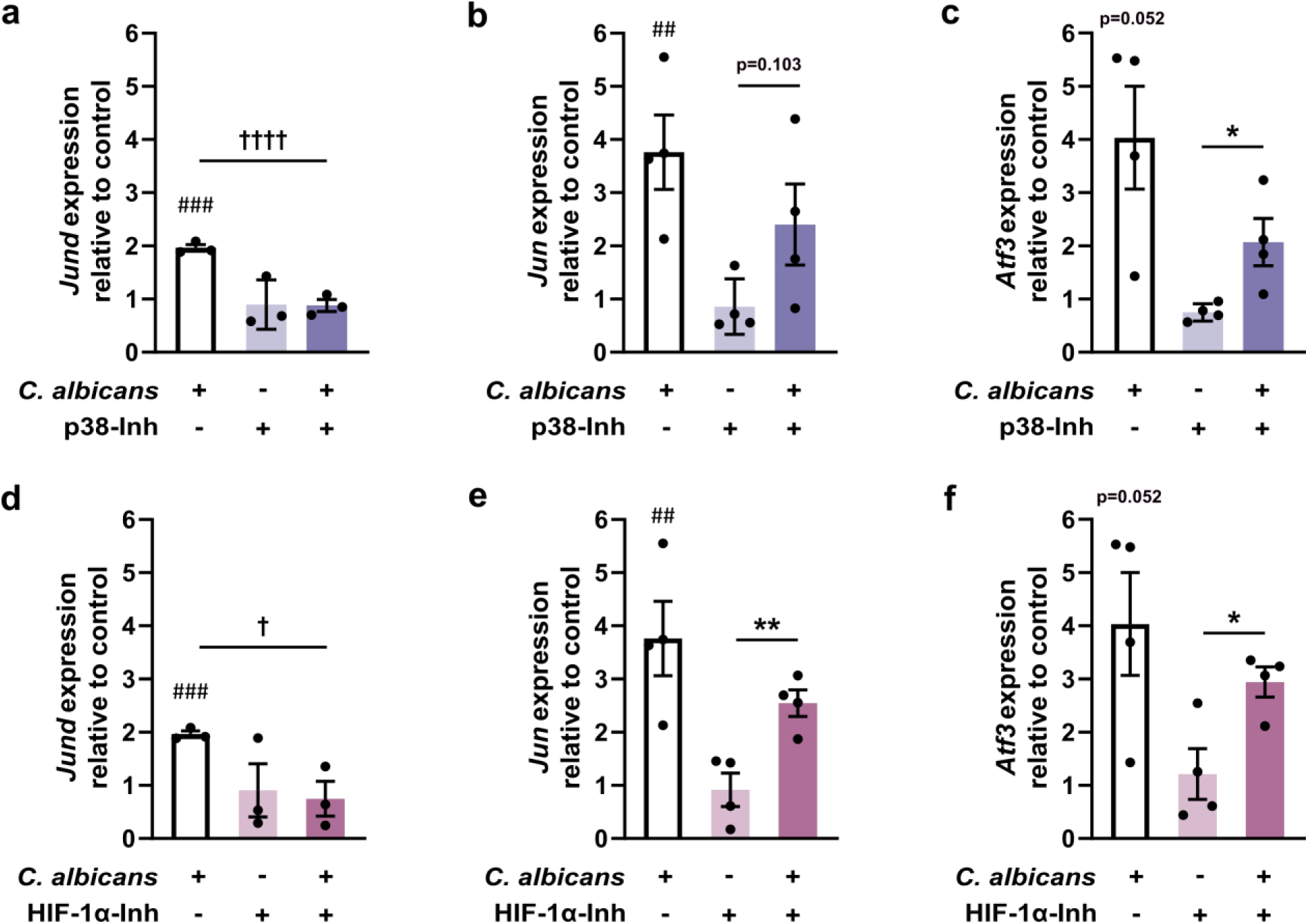
Analysis of AP-1 transcription factor activation by MOI 5 live *Candida albicans* in melanoma cells. Gene expression of *Jund*, *Jun* and *Atf3* analysis induced by six hours of stimulation with *C. albicans* alone or treated with **a**, **b**, **c** p38 inhibitor or **d**, **f**, **g** HIF-1α inhibitor. Individual values and mean ± SEM are shown. **a** and **d** (*n* = 3 biologically independent samples); **b**, **c**, **e** and **f** (*n* = 4 biologically independent samples). ^##^ and ^###^ denote *P* < 0.01 and *P* < 0.005 respect to control without *C. albicans*, respectively; * and ** denote *P* < 0.05 and *P* < 0.01 of each inhibitor compared to inhibitor with *C. albicans*, respectively; † and †††† denote *P* < 0.05 and *P* < 0.001 *C. albicans* group compared to inhibitors with *C. albicans* groups, respectively (two-tailed, unpaired, *t-student* test).

The results showed that the gene expression of all three genes and particularly *Jund* was reduced by p38-MAPK inhibition and, to a lesser extent, by HIF-1α inhibition. These results suggest that proteins encoded by these genes in addition to *Fos* may also form the AP-1 complex that activates VEGF secretion induced by *C. albicans*, and that both p38-MAPK and HIF-1α are involved in AP-1 activation, although other processes may also take part.

### 2.6. *Central carbon metabolism is altered in melanoma cells due to* C. *albicans stimulation*

To confirm the metabolic reprogramming induced by *C. albicans* in melanoma cells suggested by the transcriptomic analysis, Extracellular Acidification Rate (ECAR) analyses were performed. The results showed a significant acidification of the media after six hours when the cells were exposed to *C. albicans*, which is potentially a consequence of an increase in aerobic glycolysis (**Fig. 6a**).

**Figure 6.**
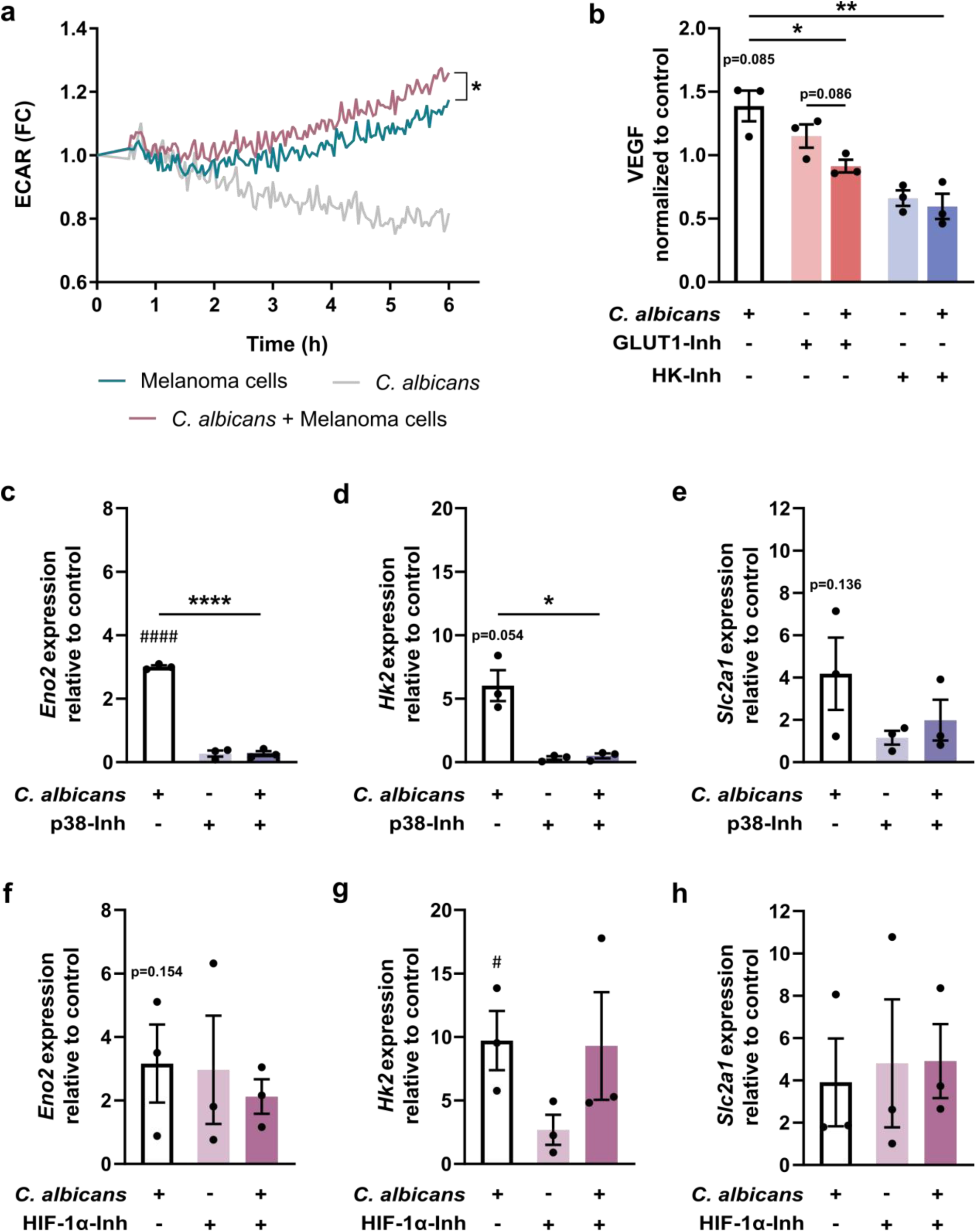
Analysis of metabolic changes induced by live *Candida albicans* in melanoma cells. **a** Extracellular acidification rate (ECAR) of melanoma cells in controls and in cell in contact with *C. albicans* for six hours. Data shown as the mean of the fold change (FC) of initial fluorescence values in three independent experiments. **b** VEGF cytokine release induced by six hours of stimulation with *C. albicans* alone or treated with glucose transporter 1 (GLUT1) inhibitor (BAY-876; 25 nM) and hexokinase (HK) inhibitor (2-DG; 5 mM). Gene expression of aerobic glycolysis related genes (*Eno2*, *Hk2* and *Slc2a1*) by six hours of stimulation with *C. albicans* alone or treated with **c**, **d**, **e** p38 inhibitor (SB 203580, 10 µM) or **f**, **g**, **h** HIF-1α inhibitor (LW6, 10 μM). For all data, individual values and mean ± SEM are shown. **a** (*n* = 4 biologically independent samples); **b**, **c**, **d**, **e**, **f** and **g** (*n* = 3 biologically independent samples). ^#^ and ^####^ denotes *P* < 0.05 and *P* < 0.001 respect to control without *C. albicans*, respectively; * and *** denote *P* < 0.05 and *P* < 0.005 of each inhibitor compared to inhibitor with *C. albicans*, respectively (two-tailed, unpaired, *t-student* test).

The interconnection between these metabolic changes and the pro-angiogenic VEGF release induced by *C. albicans* was further studied using molecular inhibitors for the glucose transporter GLUT1 (coded by *Slc2a1* gene) and the glycolytic enzyme hexokinase (HK2). Inhibition using either of the two inhibitors significantly decreased *C. albicans*-induced VEGF release in melanoma cells (**Fig. 6b**).

Finally, to determine the influence of p38-MAPK and HIF-1α on metabolic reprogramming, the expression of the genes *Eno2*, *Hk2* and *Slc2a1* was analysed after inhibition of both pathways. Inhibition of p38-MAPK resulted in a reduction in expression of all three genes, with statistically significant differences for *Eno2* and *Hk2* (**Fig. 6c-e**). Conversely, HIF-1α inhibition did not significantly inhibit gene expression of any of the three genes (**Fig. 6f-h**). These results suggest that p38-MAPK activation drives metabolic reprogramming while the HIF-1α does not.

## 3. Discussion

Microbial infections have been associated with cancer development and progression, with most of the research focussing on viruses and bacteria, while fungi remain overlooked. Despite this, an increasing number of studies have highlighted the role of fungi in cancer, in particular *C. albicans* ^5–7^. *C. albicans* is a commensal fungus often found in the human mycobiota of the skin, among other areas ^9,10^, where one of the most aggressive and deadly tumours, melanoma, develops ^31^. The aim of this study was to investigate the impact of *C. albicans* presence on cells from this type of cancer.

According to the latest “Hallmarks of Cancer” proposed by Douglas Hanahan ^32^, tumour cells acquire a range of capabilities essential for developing malignant phenotypes, such as uncontrolled proliferation and augmented invasiveness, among others. In this regard, our results showed an increase in adhesion to LSECs as well as migratory capacity of *C. albicans*-exposed melanoma cells, although no increase in their proliferation was observed. These results are in line with the results obtained by Vadovics *et al.* ^16^ for oral squamous cell carcinoma (OSCC) cell lines. Furthermore, LSECs exposed to supernatants obtained from live *C. albicans*-stimulated melanoma cells showed a significantly increased migratory capacity. This pro-angiogenic capacity could be caused by the production of growth factors and cytokines (i.e., VEGF, bFGF, IL-8, or angiopoietin) by melanoma cells ^33^.

In accordance with our *in vitro* data, pre-incubation of melanoma cells with *C. albicans* enhanced their metastatic capacity in an *in vivo* murine model of metastatic progression, increasing macro- and microscopic hepatic metastatic area. These results are in line with previous studies in which mice injected with both melanoma cells and *C. albicans* presented an augmented liver metastasis and increased TNF and IL-18 release, in comparison to mice injected only with melanoma cells ^25^. Moreover, studies with OSCC mouse models suggested that *C. albicans* can initiate cancer and contribute to its progression and development *in vivo*^16,23,24,34^.

Once we observed the phenotypical effects exerted by *C. albicans in vitro* and *in vivo*, a transcriptomic analysis of melanoma cells revealed that they respond to *C. albicans* primarily by activating the MAPK and HIF-1 signalling pathways. Exposure to *C. albicans* led to the overexpression of several MAPK-related components, including kinases, transcription factors and signalling regulators, as well as metabolic genes associated with hypoxia, such as glycolytic enzymes, glucose transporters, lactate production, and tricarboxylic acid cycle inhibitors. Remarkably, only one overexpressed gene, the pro-angiogenic VEGF, connected all these pathways, highlighting its potentially pivotal function in this response.

Subsequently, we investigated whether the activation of melanoma cells by *C. albicans* depends on fungal morphology, viability, or molecules released by the fungus. We examined the release of VEGF and gene expression of *Fos*, which was also found overexpressed in the transcriptomic data and extensively reported in the response to *C. albicans* hyphae ^28,35–37^. The highest increase in both cases was induced by live *C. albicans*, indicating that fungal viability is important for the process. This viability allows morphological changes that can increase its virulence ^38^ and to secrete proteins or toxins like candidalysin which may trigger an inflammatory response specifically linked to activation of the MAPK/c-Fos pathways ^39,40^. This is also consistent with the reduced virulence of yeast-locked *C. albicans* mutants exhibited in mice ^41^, and the lack of response of epithelial cells to HK-yeasts ^28^. Therefore, further studies were performed using live *C. albicans* yeast.

Investigation of the involvement of several receptors, mainly PRRs, already linked to *C. albicans* recognition, including TLRs, CLRs, EGFR, and EphA2 ^42,43^, in the responses described here indicated that only the inhibition of MyD88-dependent TLRs (all TLRs, except TLR3) significantly reduced VEGF release. These results support the role of TLRs not only in recognition of *C. albicans* by melanoma cells, but also in MAPK/AP-1 activation and subsequent VEGF release, as previously observed ^44^. However, our results demonstrated that while c-Fos/AP-1 inhibition reduced VEGF release, c-Fos protein expression is not essential for TLR-initiated VEGF secretion. This suggests that TLRs may activate other components of the AP-1 complex. In fact, AP-1 is a dimeric transcription factor widely related to cancer development that can be composed of different proteins from the Fos, Jun, ATF, and MAF families ^30^.

In contrast, EphA2 inhibition completely suppressed c-Fos protein expression, which is consistent with EphA2 role in activating inflammatory responses against *C. albicans* via the c-Fos/AP-1 pathway ^45^. However, no effect on VEGF release was observed. Therefore, although it cannot be ruled out that c-Fos may play a role in in *C. albicans*-induced VEGF release conforming AP-1 heterodimers with other proteins from the families abovementioned (i.e., c-Jun) ^30^, these findings suggest that AP-1 homo/heterodimers that do not include c-Fos may be enough to drive VEGF production in melanoma cells ^46–48^. Taking all this into account, we propose that *C. albicans* recognition by TLRs stimulates the activation of different AP-1 complex proteins, which subsequently leads to VEGF release. In contrast, EphA2 receptor activity appears to activate c-Fos, which needs additional components to form functional heterodimers of AP-1 to stimulate VEGF production.

Next, we analysed the effect of the two main pathways enriched in our transcriptomic analyses on VEGF release, namely the MAPK pathways (specifically the ERK1/2, JNK and p38) and HIF-1 (specifically the HIF1α subunit). In addition to them, the NF-κB pathway was included, as various cell types respond to *C. albicans* through its activation ^28,37,49,50^. Of all of them, we found that only inhibition of the p38-MAPK and HIF-1α, but not ERK1/2, JNK, or NF-κB, led to a significant reduction in VEGF release from melanoma cells stimulated with *C. albicans*. These results agree with previous studies indicating that the p38-MAPK drives VEGF production through the formation of AP-1 complexes in different cell lines ^44,51–53^, whilst HIF-1α is a well-established promoter of VEGF release ^54^. Additionally, it has been already stated that the abovementioned TLRs induce activation of p38-MAPK and c-Fos after *C. albicans* recognition ^28,55^. Moreover, Roselletti et al., (2019) showed that *C. albicans*-hyphae carriers with vaginal candidiasis exhibited enhanced expression of TLR2, TLR4 and EphA2, as well as p38-MAPK and c-Fos/AP-1 compared to *C.albicans*-yeast carriers or healthy patients. Furthermore, the activation of the p38-MAPK has also been directly associated with increased migration and proliferation of melanoma cells ^57,58^ and pancreatic cancer cells ^59^, in agreement with our results.

As expected, we confirmed that both the p38-MAPK and HIF-1α participate, at least in part, in c-Fos gene and protein expression. Further studies will explore whether these two pathways are linked in these events. Any of the three MAPK pathways (ERK1/2, JNK and p38) can lead to HIF-1α activation ^60–63^. Similarly high HIF-1α activity is able to activate MAPK signalling ^64^. This could explain the impact of HIF-1α on c-Fos expression, as already reported for c-Jun ^65^.

In our transcriptomic analysis the expression of other subunits of the AP-1 complex were found upregulated, specifically *Atf3*, *Jun*, and *Jund* genes. Further studies showed that all three genes and specifically *Jund* gene expression are regulated by both p38-MAPK and HIF-1α, probably in the same fashion as *Fos*. However, similar to c-Fos, their expression was not totally repressed by p38-MAPK or HIF-1α inhibitors, suggesting that a combination of processes are implicated in their induction. Some of these results align with those reported by Catar *et al.* ^48^ identifying the c-Fos/c-Jun AP-1 heterodimer complex as a critical regulator of VEGF release, with c-Fos playing a particularly prominent role in human vascular endothelial cells. Similar to our observations, they found that the expression of *FOS* was significantly higher than *JUN*. However, the impact of limited c-Fos or excess c-Jun on VEGF production remains to be fully elucidated ^47^.

Apart from VEGF release, transcriptomic results indicated an upregulation of genes associated with aerobic glycolysis, suggesting that melanoma cells shift their metabolism towards this pathway in response to the fungus. To confirm this metabolic reprograming, an ECAR assay was performed, which showed an increased acidification rate of the media as a result of lactate produced from pyruvate. Increased aerobic glycolysis has been often associated with the high energy demands that cancer cells present to support their migration, invasion, and proliferation ^66–69^, but it has been also described in response to *C. albicans* in both non-tumour ^70–72^ and tumour ^16^ cell lines. Moreover, this process can be dependent on several mechanisms, including HIF-1α and MAPKs ^66,68,69^.

Therefore, we studied the relevance of this metabolic reprograming in the release of VEGF stimulated by *C. albicans* on melanoma cells. Data showed a decrease in VEGF release when inhibiting the glycolytic enzyme HK2 and GLUT1, suggesting that the glycolytic pathway is necessary to maintain this response. Indeed, lactate, the end-product of aerobic glycolysis, is reported to play a key role in VEGF production ^73–76^, and HIF-1α accumulation ^77^. Moreover, it has been described that increasing lactate production promotes resistance of cancer cells to chemotherapy ^78^.

Finally, we investigated the role of the p38-MAPK and HIF-1α on the expression of the genes involved in aerobic glycolysis *Hk2*, *Slc2a1*, and *Eno2*. Our results revealed that p38-MAPK upregulates *Eno2* and *Hk2* expression and *Slc2a1* to a lesser extent. Indeed, the involvement of p38-MAPK in the shift from oxidative phosphorylation to aerobic glycolysis has been observed in sepsis ^79^ and in pancreatic cancer cells ^59^. However, although HIF-1α is frequently associated with the metabolic reprogramming in cancer cells ^80^, we did not observe it in this case, and therefore, further studies will be needed to clarify this point.

In conclusion, this study shows that *C. albicans* infection induces changes at the gene expression level in tumour cells, resulting in a more aggressive phenotype and the formation of a tumour microenvironment that would favour both the development and progression of cancer (**Fig. 7**). The fungus promotes melanoma cell migration, adhesion to LSECs, metastatic capacity, release of the pro-angiogenic VEGF, and metabolic reprogramming. Our data supports a model in which MyD88-mediated TLRs and EphA2 recognize *C. albicans* and induce VEGF release and c-Fos expression, respectively. Both p38-MAPK and HIF-1α are activated, contributing to VEGF release via activation of c-Fos/AP-1 and possibly other AP-1 subunits, with p38-MAPK being responsible for the metabolic reprogramming. This, in turn, supports the maintenance of a pro-angiogenic microenvironment. Further studies will elucidate the mechanisms underlying the pro-tumour and pro-metastatic processes observed and potentially open the possibility of using antifungal therapies as complements to traditional treatments for melanoma.

**Fig. 7.**
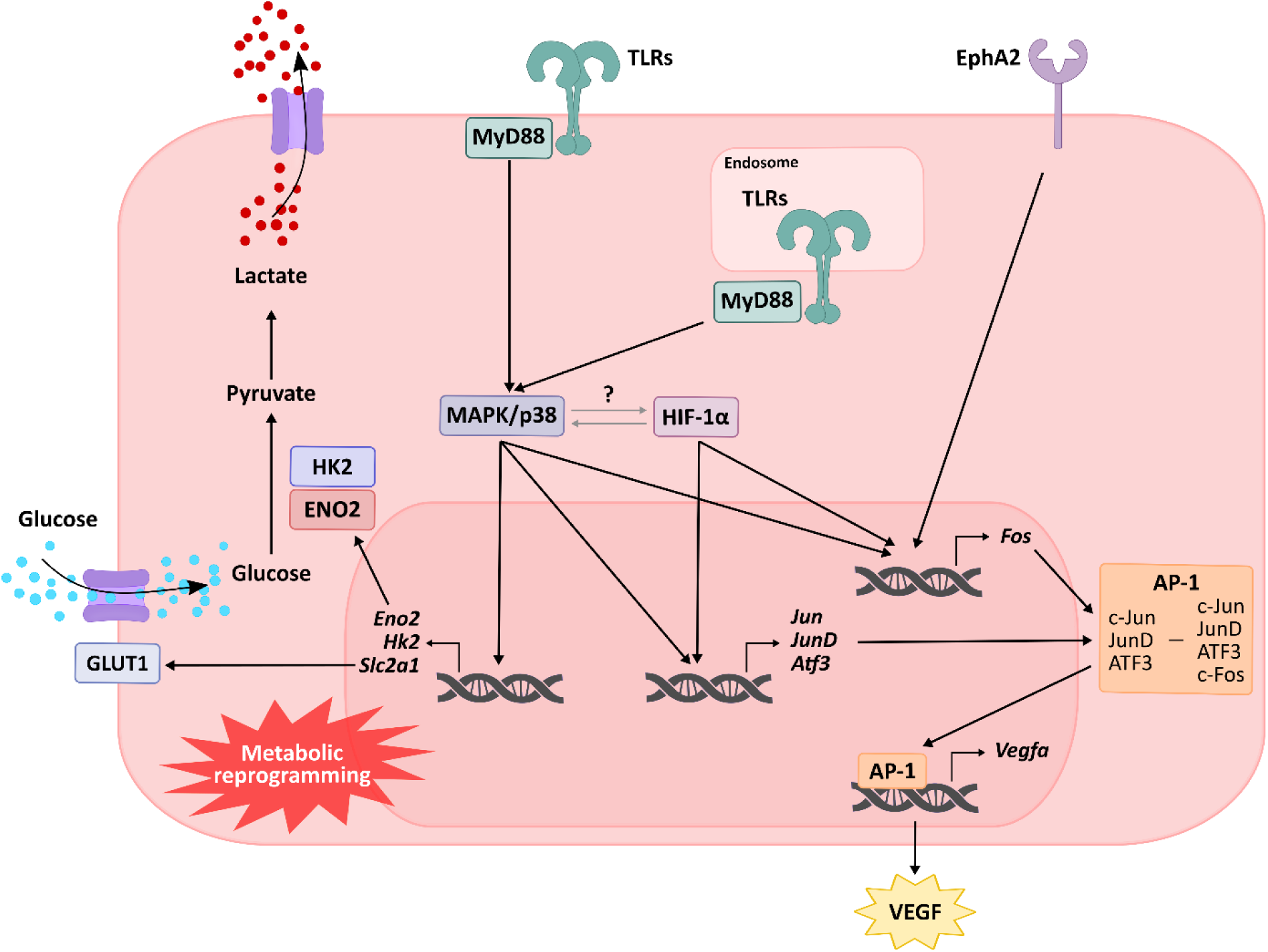
Graphical representation of the hypothesized melanoma response to *C. albicans*. *C. albicans* is recognized by TLRs and EphA2, triggering VEGF release and c-Fos expression, respectively. Intracellularly, the p38 and HIF-1α pathways are activated, promoting VEGF release through the activation of c-Fos/AP-1 and potentially other AP-1 subunits. Additionally, the p38 pathway plays a key role in driving metabolic reprogramming.

## 4. Materials and methods

### 4.1. Microorganisms

*Candida albicans* CECT 13062 strain (Spanish Type Culture Collection) isolated from a patient with systemic candidiasis was used with the approval of the Ethics Committee of the University of the Basque Country (UPV/EHU; M30/2019/114). The fungus was grown in Sabouraud Dextrose Agar (SDA; Panreac Química SLU, Castellar del Vallès, Spain) at 37°C overnight. One day prior to the experiment, 10^5^ cells/mL were inoculated in Sabouraud Dextrose Broth (SDB; Panreac Química SLU) and incubated at 37°C and 120 rpm overnight. The day of the experiment, yeasts were collected by centrifugation and washed twice with Dulbecco’s Phosphate Buffer Saline (DPBS; Corning, VWR, PA, USA). Fungal cell numbers were calculated for each experiment using a hemocytometer.

All the experiments were carried out with live *C. albicans* unless otherwise specified. In order to obtain the inactive form of *C. albicans*, either yeast or hyphae, the fungus was heat-killed (HK-*C. albicans*) adjusting its concentration to 10^7^ cells/mL in DPBS and incubating for 30 minutes at 70°C. Conditioned media was obtained by seeding 5 × 10^5^ melanoma cells/well in 12-well plates, stimulating them with live *C. albicans* at a multiplicity of infection (MOI; *C. albicans*:melanoma cells) of 5 for six hours, and then collecting the supernatants, which were stored at –20°C until further use.

### 4.2. Cells and culture conditions

B16-F10 melanoma cells (ATCC CRL-6475; VA, USA) and immortalized murine liver sinusoidal endothelial cells (LSECs; Innoprot, Spain) were kept in Roswell Park Memorial Institute (RPMI) 1640 medium with 10% heat-inactivated fetal bovine serum (FBS), 2 mM L-glutamine, 100 U/mL penicillin, 100 µg/mL streptomycin and 0.25 μg/mL amphotericin B (all from Sigma-Aldrich, MO, USA), in a 5% CO_2_ atmosphere at 37°C.

### 4.3. Wound healing assay

To evaluate cell migration, 1.5 x 10^5^ melanoma cells/well were seeded and cultured overnight in 24-well plates (Sarstedt, Germany) for obtaining a cell monolayer. The following day, a vertical and a horizontal scratch was made using a 1,000 µL pipette tip followed by two washing steps twice with DPBS to remove detached cells. Then, fresh media alone or with MOI 0.25 and 0.5 of HK-*C. albicans* yeast was added to each well. A Nikon Eclipse TE2000-U inverted microscope (Nikon, Minato, Tokyo, Japan) was utilized to photograph the same position of scratches at 0 and 24 hours after scratching. The percentage of scratch reduction (i.e., wound closure) was calculated as follows; *Wound closure (%) = [(Initial wound area – Final wound area)/Initial wound area] x 100*, and normalized to the untreated control.

### 4.4. Melanoma cells adhesion to LSECs assay

To study the effect of *C. albicans* on the adhesion capacity of melanoma cells to endothelial cells (LSECs), the day before the experiment 2.5 × 10^5^ LSECs/well were seeded in a 48-well plate (Costar, WA, USA) and incubated overnight. Additionally, 1 × 10^6^ melanoma cells were seeded in 25 cm^2^ cell culture flasks (Sarstedt) with or without HK-*C. albicans* yeast at MOI 10. The following day, melanoma cells were harvested and stained using 1 µM of carboxyfluorescein succinimidyl ester (CFSE) (Invitrogen, Carlsbad, CA USA) per 1 x 10^6^ cells in DPBS for 30 min, followed by washing in RPMI supplemented with 10% of FBS. Then, 2.5 × 10^4^ melanoma cells per well were added to the LSECs monolayers. The resulting co-cultures were maintained for 10 min at 37°C and total emitted fluorescence was measured using Synergy HT (BioTek, USA) with the Gen5 program (BioTek) and defined as initial fluorescence. Then, co-cultures were extensively but delicately washed with culture medium to prevent removing of adherent cells. The fluorescence emitted by adhered cells was again measured (final fluorescence). Finally, the percentage of adhered cells was calculated by the subtraction of background fluorescence (fluorescence emitted by LSECs alone) as follows: *Cell adhesion (%) = [(Final fluorescence – Background fluorescence)/(Initial fluorescence - Background fluorescence)] x 100*.

### 4.5. Angiogenesis promotion assay

The pro-angiogenic microenvironment promotion was determined by seeding in 12-well plates 5 x 10^5^ melanoma cells/well with an MOI 5 of live *C. albicans* for six hours. The collected supernatant was then centrifuged to discard tumour and fungal debris, and the obtained conditioned media was stored at 4°C with 0.25 μg/mL amphotericin B until the following day. In parallel, 1.25 x 10^5^ LSECs/well were seeded in 24-well cell culture plates overnight for obtaining cell monolayers. The following day, wound healing assays were performed as described above using the conditioned media stored as culture media and measuring wound closure percentage of LSECs after eight hours of exposure.

### 4.6. Cell proliferation assay

Cell proliferation assays by flow cytometry were performed in the Advanced Research Facilities from the UPV/EHU (SGIker) using a Beckman Coulter Gallios cytometer (Beckman Coulter, CA, USA).

Melanoma cells were first stained with the fluorescent dye CFSE as before. They were then resuspended in RPMI supplemented with 10% FBS and seeded at 2.5 × 10^5^ cells/well in 6-well plates. At 0 hours, the cells were analysed by flow cytometry using unstained cells as controls. The rest of the cells were exposed to fresh media alone or with MOI 1 and 10 of HK-yeast and after 24, 48 and 72 hours, the cells were detached, resuspended in DPBS and analysed in the same way. The flow cytometer was set up to analyse 10,000 events from each sample, exciting the sample at 492 nm and measuring the emitted fluorescence at 517 nm. Results were analysed using Flowing Software 2 (https://bioscience.fi/services/cell-imaging/flowing-software/; Turku Bioscience Centre, Turku, Finland).

### 4.7. In vivo *development of hepatic metastasis*

The effect of *C. albicans* on the metastatic capacity of melanoma cells *in vivo* was studied using a murine hepatic metastasis model. All the *in vivo* experiments were performed in the Animal Unit Service of the SGIker and were approved by the Animal Experimentation Ethics Committee from the UPV/EHU (M20/2022/052). Syngeneic C57BL/6J mice (male, 6-8 weeks old) purchased from Janvier Labs (Saint-Berthevin, France) were used.

First, 1 × 10^6^ melanoma cells were seeded in 25 cm^2^ cell culture flasks and incubated overnight. The following day, live *C. albicans* was added at MOI of 1 to the *C. albicans*-stimulated melanoma group. After six hours of co-incubation, melanoma cells of control and *C. albicans*-stimulated groups were detached from the flasks and adjusted to 1 x 10^6^ cell/mL in DPBS supplemented with 25 μg/mL of amphotericin B in order to chemically inactivate the remaining *C. albicans*.

For the establishment of the hepatic metastasis, mice were first anesthetized, and then, 1 × 10^5^ melanoma cells/100 µl of DPBS per mouse, either previously stimulated with *C. albicans* or not, were injected intrasplenically. Mice were anesthetized and sacrificed 14 days after tumour cells injection, the liver was collected, fixed and paraffin-embedded for histological analyses after hematoxylin-eosin (H&E) staining. The macro- and microscopic regions of the liver occupied by tumour cells (tumour area) were quantified in the extracted organ and in 7 μm-thick tissue sections, respectively. Three different sections of each liver were evaluated allowing 500 μm separation between each of them. This assay was performed twice with at least five mice per group.

### 4.8. RNA-seq and bioinformatic analysis

For transcriptomic analysis, 1 × 10^6^ melanoma cells/well were seeded in 6-well plates (Sarstedt), stimulated with MOI 1 of live *C. albicans* and after six hours total RNA isolation was performed by Total RNA Isolation Kit (NzyTech), following manufacturer’s instructions. Total RNA quantity was measured using the Qubit RNA Assay Kit (Thermo Fisher Scientific) and its quality integrity was measured examined by RNA Nano Chips in a 2100 Bioanalyzer (Agilent Technologies, Santa Clara, CA) in the Center for Cooperative Research in Biosciences (CICBiogune) in Derio, Spain.

Then, RNA libraries were prepared with the TruSeq® Stranded mRNA Library Prep” kit (Illumina Inc. Cat. # 20020594) and TruSeq RNA Single Indexes (Illumina Inc. Cat. # 20020492 and 20020493), Truseq RNA Sample Preparation Kit v2 (Illumina, CA, USA) and sequenced on HiScanSQ NovaSeq6000 Platform (Illumina Inc.) to generate 50 100 base single paired-end reads. The quality control of raw sequence data was performed using FastQC v.0.11.9.

Using STAR software ^81^, FASTQ files were aligned to Mouse genome data base mm10 and read counts obtained for each gene. Differential expression analysis was performed with the DESeq2 (data available through the Gene Expression Omnibus (https://www.ncbi.nlm.nih.gov/geo/ under accession number GSE285124).

The differentially expressed genes (DEGs), Gene Ontology (GO) and KEGG pathway enrichment analysis were determined using the online iDEP 1.0 tool (http://149.165.154.220/idep10/, BioMed Central Ltd, BCM Informatics) and Deseq2 method. For all of them the cut-off was assigned above the │FC│ > 1.5 and a p_adj_ < 0.05. Further protein interaction networks studies were performed using the STRING tool. Furthermore, a second bioinformatics analysis of the results considering only the p_adj_ < 0.05 was also done.

### 4.9. Measurement of Vascular Endothelial Growth Factor (VEGF) release

For cytokine measurement, 5 × 10^5^ melanoma cells/well were seeded in 12-well plates (Sarstedt) and stimulated with an MOI 5 of live *C. albicans*. After six hours the supernatants were collected and stored at –20°C until use. Enzyme-linked Immunosorbent Assays (ELISA) to detect Vascular Endothelial Growth Factor (VEGF) was performed using commercial VEGF-165 Mouse ELISA Development Kit (ABTS) (PeproTech® by Invitrogen, Thermo Fisher Scientific).

### 4.10. Gene expression measurement by RT-qPCR analysis

For RT-qPCR, 1 × 10^6^ melanoma cells/well were seeded in 6-well plates, stimulated with an MOI of 5 of live *C. albicans* and after six hours total RNA was isolated and measured using the Nanodrop ND 1000 (Thermo Fisher Scientific) before being stored at –80°C until used. cDNA was synthesized using the NZY First-Strand cDNA Synthesis Kit (NzyTech). RT-qPCR was performed using the NZYSupreme qPCR Green Master Mix (2x) (NzyTech) in a CFX96 Touch Real-Time PCR Detection System (Bio-Rad, CA, USA) thermal cycler. The primers used are listed in **Supp. Table 3**. To determine the relative concentration of mRNAs relative to control the threshold cycle (Ct) values were normalized to the *Rpl19* housekeeping gene using the ΔΔCt method.

### 4.11. Electrophoresis and Western blotting

First, 5 × 10^5^ melanoma cells/well in 12-well plates were seeded and stimulated for six hours with an MOI of 5 of live *C. albicans*. Protein extraction was performed using RIPA lysis buffer (Sigma-Aldrich) containing 1 mM protease inhibitors (Phenylmethylsulfonyl fluoride, PMSF, Sigma-Aldrich) and incubated on ice for 10 min. Each well was then scrapped, collected, and incubated on ice another 10 min. Finally, the samples were centrifuged at maximum speed for 5 min and the supernatants stored at -20°C until use.

For Western Blotting (WB), samples were loaded into 10% SDS-polyacrylamide gels and run at 70 mA, 100 W and 200 V for 45 min in a Miniprotean II (Bio-Rad), using the Page Ruler Plus protein standard (Thermo Fisher Scientific). Proteins were then transferred to Hybond-P PVDF membranes (GE Healthcare) for 1 h at 400 mA. Membranes were then blocked for 2 h with Tris Buffered Saline Milk (TBSM; 50 mM tris-HCl pH 7.5, 150 mM NaCl, 0.1% (v/v) Tween 20 and 5% (w/v) low fat dried milk, both from Sigma-Aldrich, MO, USA). Rabbit monoclonal antibodies against c-Fos (YA506, HY-P80081) and the housekeeping protein β-Tubulin (HY-P80487) diluted 1:1,000 in TBSM were used as primary antibodies (MedChemExpress, NJ, USA) and incubated with the membrane overnight at 4°C. After three 5 min washes with TBS, goat anti-rabbit IgG (Invitrogen, Thermo Fisher Scientific) diluted 1:5,000 in TBSM was used as secondary antibody and incubated with the membrane for 30 min. All the steps were carried out at room temperature and agitation unless otherwise indicated. Membranes were developed using the NZY Standard ECL (NzyTech) in a G:BOX Chemi system (Syngene). Densitometry analysis of WBs was performed using Image J (Bethesda, MD, USA).

### 4.12. Inhibition of signalling and metabolic pathways

To inhibit different metabolic and signalling pathways, melanoma cells were treated with inhibitors for one hour at 37°C and 5% CO_2_ before *C. albicans* was added. The inhibitors of hexokinase (2-deoxyglucose, 2-DG; 5 mM), GLUT1 (BAY 876; 25 nM), SYK (Piceatannol, 30 µM), MyD88 (TJ-M2010-5, 30 µM), EGFR (PD153035, 1 µM), EphA2 (NVP-BHG712, 1 µM), ERK1/2 (PD-0325901; 1 µM), JNK (SP600125, 10 µM), p38a (SB 203580, 10 µM), NF-κB (BAY 11-7082, 2 µM), c-Fos/AP-1 (T-5224, 10 µM) and HIF-1α (LW6, 10 μM) were obtained from MedChemExpress.

### 4.13. Extracellular Acidification Rate (ECAR) assay

Glycolytic activity was assessed using the Extracellular Acidification Rate (ECAR) as a proxy, using the Agilent MitoXpress® pH-Xtra kit (Agilent, CA, USA) following the manufacturer’s instructions.

First, 2.5 x 10^4^ melanoma cells were seeded per well in black 96-well plates with clear bottoms (Costar, Corning, NY, USA) and incubated overnight. The cells were washed twice with 100 μL of pre-warmed respiration buffer adjusted to pH 7.4. After, 90 μL of fresh respiration buffer was added to each well and immediately cells were infected with an MOI of 5 of live *C. albicans*. Finally, 10 μL of pH-Xtra reagent was added to each well and the plates were incubated in the FlexStation 3 CO_2_ free machine (Molecular Devices, CA, USA) for six hours at 37°C, exciting the sample at 380 nm and measuring the fluorescence emitted by pH-Xtra reagent at 615 nm.

### 4.14. Statistical analysis

Statistical analysis was performed using the IBM SPSS statistical software (version 28.0.1.1; Professional Statistic, IL, USA). All the experiments were performed at least three times and analysed using two-tailed unpaired *t-student* test. The criterion for significance was set at p < 0.05 for all comparisons.

## Acknowledgments

This research was funded by the Basque Government, grant numbers IT1362-19 and IT1657-22. L.M-S, M.A, O.R and L.A-D have received a predoctoral grant from the Basque Government and L.A-F from the University of the Basque Country (UPV/EHU). AP was supported by a fellowship from the Basque Government Postdoctoral Program. CIC bioGUNE support was provided by the Basque Department of Industry, Tourism and Trade (Etortek, Elkartek and Emaitek Programs), the Innovation Technology Department of Bizkaia County, CIBERehd Network, and Spanish MINECO the Severo Ochoa Excellence Accreditation (CEX2021-001136-S).

