## Supplementary material for "*Candida albicans* enhances melanoma cell aggressiveness through p38-MAPK and HIF-1α pathways and metabolic reprogramming": Suplemmentary data

### Supplementary data

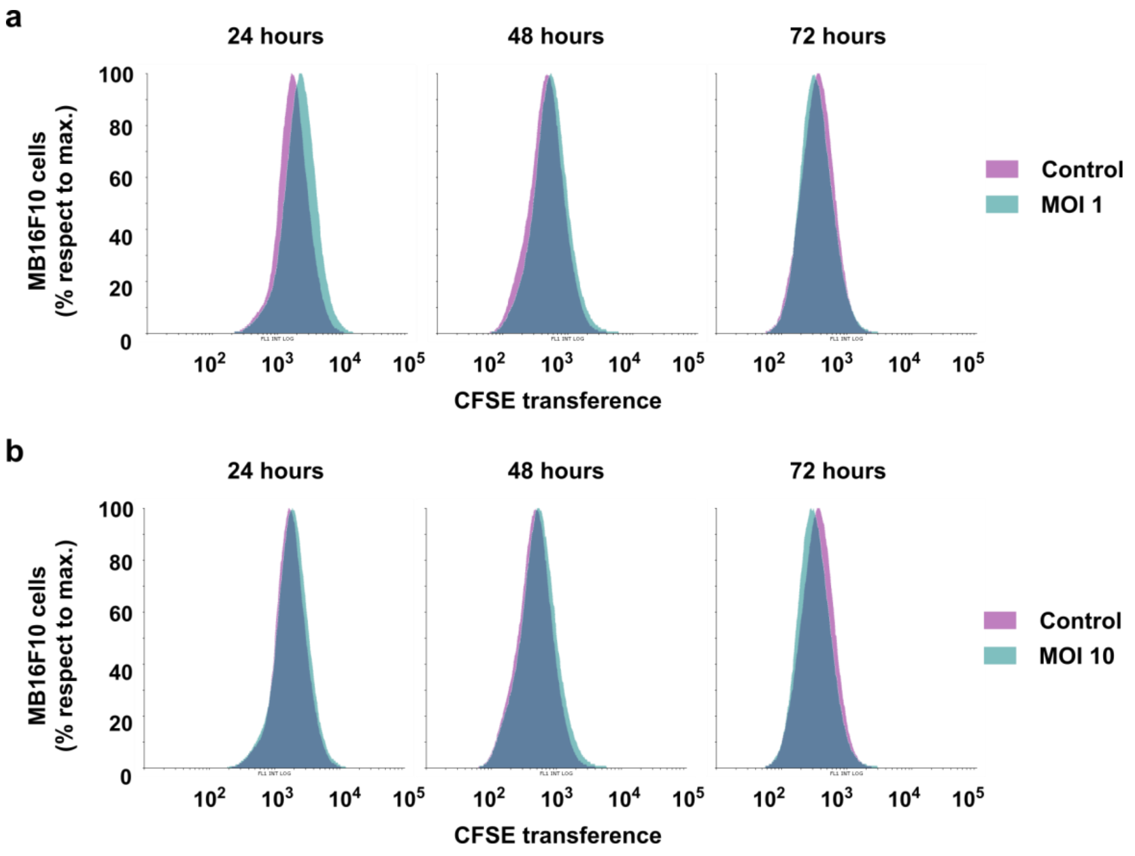


**Supplementary Figure 1. HK-*Candida albicans* effect in melanoma cells proliferation capacity.** Overlay of the fluorescence intensity of CFSE after 24, 48 and 72 hours of control and melanoma cells exposed to HK-*C. albicans* yeast at **a** MOI 1 and **b** MOI 10. In CFSE terms, more fluorescence means less cell proliferation. Representative results of three independent proliferation assays (*n* = 3 biologically independent samples) are shown.

**Supplementary Table 1. Data of gene expression of melanoma cells stimulated with live *C. albicans* MOI 1 for six hours compared to unstimulated samples.** Analysis were performed for |FC| > 1.5 and $p_{adj} \leq 0.05$ (iDEP 1.0 tool and Deseq2 method) (*n* = 3 biologically independent samples).

| Gene | Entrez Gene Name | FC | $Log_2FC$ | $p_{adj}$ value |
|---|---|---|---|---|
| ***Atf3*** | activating transcription factor 3 | 3,29 | 1,72 | 1,5E-08 |
| ***Chac1*** | cation transport regulator 1 | 2,90 | 1,54 | 6,2E-50 |
| ***Adm*** | adrenomedullin | 2,84 | 1,50 | 1,6E-02 |
| ***Ankrd37*** | ankyrin repeat domain 37 | 2,82 | 1,49 | 4,4E-09 |
| ***1600014C23Rik*** | RIKEN cDNA 1600014C23 gene | 2,69 | 1,43 | 1,3E-02 |
| ***Dusp8*** | dual specificity phosphatase 8 | 2,68 | 1,42 | 8,1E-30 |
| ***Stc1*** | stanniocalcin 1 | 2,47 | 1,31 | 6,7E-07 |
| ***Ddit4*** | DNA-damage-inducible transcript 4 | 2,30 | 1,20 | 9,7E-11 |
| ***Ier3*** | immediate early response 3 | 2,21 | 1,14 | 2,2E-04 |
| ***Bnip3*** | BCL2/adenovirus E1B interacting protein 3 | 2,19 | 1,13 | 4,8E-07 |
| ***Gm6532*** | predicted pseudogene 6532 | 2,18 | 1,13 | 1,4E-03 |
| ***Rasd2*** | RASD family, member 2 | 2,13 | 1,09 | 9,4E-03 |
| ***Trib3*** | tribbles pseudokinase 3 | 2,11 | 1,08 | 1,3E-14 |
| ***Nupr1*** | nuclear protein transcription regulator 1 | 2,04 | 1,03 | 1,6E-09 |
| ***Mpp2*** | membrane protein, palmitoylated 2 (MAGUK p55 subfamily member 2) | 1,99 | 0,99 | 1,1E-04 |
| ***Jun*** | jun proto-oncogene | 1,99 | 0,99 | 4,9E-41 |
| ***Fos*** | FBJ osteosarcoma oncogene | 1,98 | 0,98 | 5,0E-06 |
| ***Gdf15*** | growth differentiation factor 15 | 1,95 | 0,96 | 1,2E-13 |
| ***Hilpda*** | hypoxia inducible lipid droplet associated | 1,94 | 0,96 | 4,8E-07 |
| ***Gm12404*** | predicted gene 12404 | 1,93 | 0,95 | 4,6E-02 |
| ***Vegfa*** | vascular endothelial growth factor A | 1,90 | 0,93 | 1,1E-06 |
| ***Hk2*** | hexokinase 2 | 1,88 | 0,91 | 1,6E-05 |
| ***Kbtbd11*** | kelch repeat and BTB (POZ) domain containing 11 | 1,88 | 0,91 | 2,0E-06 |
| ***Rgs11*** | regulator of G-protein signaling 11 | 1,88 | 0,91 | 4,9E-02 |
| ***Plekha2*** | pleckstrin homology domain-containing, family A (phosphoinositide binding specific) member 2 | 1,86 | 0,89 | 2,8E-06 |
| ***Abhd18*** | abhydrolase domain containing 18 | 1,82 | 0,87 | 1,9E-14 |
| ***Maff*** | v-maf musculoaponeurotic fibrosarcoma oncogene family, protein F (avian) | 1,80 | 0,85 | 5,3E-03 |
| ***Ccn1*** | cellular communication network factor 1 | 1,79 | 0,84 | 3,4E-03 |
| ***Fosl2*** | fos-like antigen 2 | 1,76 | 0,82 | 6,8E-14 |
| ***Aire*** | autoimmune regulator | 1,76 | 0,82 | 1,1E-03 |
| ***Dusp1*** | dual specificity phosphatase 1 | 1,72 | 0,78 | 1,4E-05 |

| Gene | Entrez Gene Name | FC | $Log_2FC$ | $p_{adj}$ value |
|---|---|---|---|---|
| ***Epm2a*** | epilepsy, progressive myoclonic epilepsy, type 2 gene alpha | 1,72 | 0,78 | 8,1E-06 |
| ***Nppb*** | natriuretic peptide type B | 1,71 | 0,78 | 1,8E-12 |
| ***Bhlhe40*** | basic helix-loop-helix family, member e40 | 1,71 | 0,77 | 4,1E-03 |
| ***Ccng2*** | cyclin G2 | 1,70 | 0,76 | 4,3E-03 |
| ***Ddit3*** | DNA-damage inducible transcript 3 | 1,69 | 0,76 | 1,0E-21 |
| ***Ak4*** | adenylate kinase 4 | 1,69 | 0,76 | 3,1E-06 |
| ***Ero1a*** | endoplasmic reticulum oxidoreductase 1 alpha | 1,68 | 0,75 | 2,0E-04 |
| ***Ak3l2-ps*** | adenylate kinase 3-like 2, pseudogene | 1,68 | 0,75 | 8,5E-05 |
| ***Slc1a4*** | solute carrier family 1 (glutamate/neutral amino acid transporter), member 4 | 1,67 | 0,74 | 4,5E-14 |
| ***Selenbp1*** | selenium binding protein 1 | 1,66 | 0,73 | 3,6E-03 |
| ***P4ha2*** | procollagen-proline, 2-oxoglutarate 4-dioxygenase (proline 4-hydroxylase), alpha II polypeptide | 1,66 | 0,73 | 8,3E-31 |
| ***2500002B13Rik*** | RIKEN cDNA 2500002B13 gene | 1,64 | 0,72 | 3,4E-03 |
| ***Egln3*** | egl-9 family hypoxia-inducible factor 3 | 1,63 | 0,71 | 9,3E-03 |
| ***Dusp10*** | dual specificity phosphatase 10 | 1,63 | 0,70 | 5,0E-03 |
| ***Nfil3*** | nuclear factor, interleukin 3, regulated | 1,63 | 0,70 | 7,2E-08 |
| ***Slc16a3*** | solute carrier family 16 (monocarboxylic acid transporters), member 3 | 1,60 | 0,68 | 1,5E-06 |
| ***Gys1*** | glycogen synthase 1, muscle | 1,60 | 0,68 | 4,4E-06 |
| ***Pkp2*** | plakophilin 2 | 1,59 | 0,67 | 2,9E-05 |
| ***Sesn2*** | sestrin 2 | 1,58 | 0,66 | 1,0E-22 |
| ***Hoxb9*** | homeobox B9 | 1,58 | 0,66 | 1,9E-10 |
| ***Btg2*** | BTG anti-proliferation factor 2 | 1,58 | 0,66 | 2,4E-04 |
| ***Sap30*** | sin3 associated polypeptide | 1,57 | 0,65 | 7,7E-10 |
| ***Pdk1*** | pyruvate dehydrogenase kinase, isoenzyme 1 | 1,57 | 0,65 | 3,1E-03 |
| ***Rnf19a*** | ring finger protein 19A | 1,55 | 0,64 | 3,2E-06 |
| ***Rora*** | RAR-related orphan receptor alpha | 1,55 | 0,63 | 1,3E-07 |
| ***Rhob*** | ras homolog family member B | 1,55 | 0,63 | 7,8E-15 |
| ***Ppp1r15a*** | protein phosphatase 1, regulatory subunit 15A | 1,54 | 0,63 | 1,3E-11 |
| ***Slc2a1*** | solute carrier family 2 (facilitated glucose transporter), member 1 | 1,53 | 0,62 | 1,6E-07 |
| ***Plk3*** | polo like kinase 3 | 1,53 | 0,61 | 3,4E-06 |
| ***Arid5a*** | AT-rich interaction domain 5A | 1,51 | 0,60 | 5,5E-09 |
| ***Egln1*** | egl-9 family hypoxia-inducible factor 1 | 1,51 | 0,60 | 1,2E-06 |
| ***Jund*** | jun D proto-oncogene | 1,51 | 0,59 | 2,3E-05 |

**Supplementary Table 2. Data of upregulated genes expression of melanoma cells stimulated with live *C. albicans* MOI 1 for six hours compared to unstimulated samples.** Analysis were performed for $p_{adj} \leq 0.05$ (iDEP 1.0 tool and Deseq2 method) ($n$ = 3 biologically independent samples).

| Gene | Entrez Gene Name | FC | $Log_2FC$ | $p_{adj}$ value |
|---|---|---|---|---|
| ***Atf3*** | activating transcription factor 3 | 3,29 | 1,72 | 1,46E-08 |
| ***Chac1*** | cation transport regulator 1 | 2,90 | 1,54 | 6,18E-50 |
| ***Adm*** | adrenomedullin | 2,84 | 1,50 | 1,65E-02 |
| ***Ankrd37*** | ankyrin repeat domain 37 | 2,82 | 1,49 | 4,44E-09 |
| ***1600014C23Rik*** | RIKEN cDNA 1600014C23 gene | 2,69 | 1,43 | 1,34E-02 |
| ***Dusp8*** | dual specificity phosphatase 8 | 2,68 | 1,42 | 8,08E-30 |
| ***Stc1*** | stanniocalcin 1 | 2,47 | 1,31 | 6,65E-07 |
| ***Ddit4*** | DNA-damage-inducible transcript 4 | 2,30 | 1,20 | 9,74E-11 |
| ***Ier3*** | immediate early response 3 | 2,21 | 1,14 | 2,16E-04 |
| ***Bnip3*** | BCL2/adenovirus E1B interacting protein 3 | 2,19 | 1,13 | 4,75E-07 |
| ***Gm6532*** | predicted pseudogene 6532 | 2,18 | 1,13 | 1,37E-03 |
| ***Rasd2*** | RASD family, member 2 | 2,13 | 1,09 | 9,44E-03 |
| ***Trib3*** | tribbles pseudokinase 3 | 2,11 | 1,08 | 1,29E-14 |
| ***Nupr1*** | nuclear protein transcription regulator 1 | 2,04 | 1,03 | 1,57E-09 |
| ***Mpp2*** | membrane protein, palmitoylated 2 (MAGUK p55 subfamily member 2) | 1,99 | 0,99 | 1,13E-04 |
| ***Jun*** | jun proto-oncogene | 1,99 | 0,99 | 4,93E-41 |
| ***Fos*** | FBJ osteosarcoma oncogene | 1,98 | 0,98 | 4,95E-06 |
| ***Gdf15*** | growth differentiation factor 15 | 1,95 | 0,96 | 1,16E-13 |
| ***Hilpda*** | hypoxia inducible lipid droplet associated | 1,94 | 0,96 | 4,79E-07 |
| ***Gm12404*** | predicted gene 12404 | 1,93 | 0,95 | 4,55E-02 |
| ***Vegfa*** | vascular endothelial growth factor A | 1,90 | 0,93 | 1,12E-06 |
| ***Hk2*** | hexokinase 2 | 1,88 | 0,91 | 1,56E-05 |
| ***Kbtbd11*** | kelch repeat and BTB (POZ) domain containing 11 | 1,88 | 0,91 | 2,01E-06 |
| ***Rgs11*** | regulator of G-protein signaling 11 | 1,88 | 0,91 | 4,88E-02 |
| ***Plekha2*** | pleckstrin homology domain-containing, family A (phosphoinositide binding specific) member 2 | 1,86 | 0,89 | 2,82E-06 |
| ***Abhd18*** | abhydrolase domain containing 18 | 1,82 | 0,87 | 1,86E-14 |
| ***Maff*** | v-maf musculoaponeurotic fibrosarcoma oncogene family, protein F (avian) | 1,80 | 0,85 | 5,25E-03 |
| ***Ccn1*** | cellular communication network factor 1 | 1,79 | 0,84 | 3,37E-03 |
| ***Fosl2*** | fos-like antigen 2 | 1,76 | 0,82 | 6,81E-14 |
| ***Aire*** | autoimmune regulator | 1,76 | 0,82 | 1,07E-03 |
| ***Dusp1*** | dual specificity phosphatase 1 | 1,72 | 0,78 | 1,45E-05 |

| Gene | Entrez Gene Name | FC | $Log_2FC$ | $p_{adj}$ value |
|---|---|---|---|---|
| ***Epm2a*** | epilepsy, progressive myoclonic epilepsy, type 2 gene alpha | 1,72 | 0,78 | 8,10E-06 |
| ***Nppb*** | natriuretic peptide type B | 1,71 | 0,78 | 1,81E-12 |
| ***Bhlhe40*** | basic helix-loop-helix family, member e40 | 1,71 | 0,77 | 4,13E-03 |
| ***Ccng2*** | cyclin G2 | 1,70 | 0,76 | 4,29E-03 |
| ***Ddit3*** | DNA-damage inducible transcript 3 | 1,69 | 0,76 | 1,01E-21 |
| ***Ak4*** | adenylate kinase 4 | 1,69 | 0,76 | 3,14E-06 |
| ***Ero1a*** | endoplasmic reticulum oxidoreductase 1 alpha | 1,68 | 0,75 | 2,00E-04 |
| ***Ak3l2-ps*** | adenylate kinase 3-like 2, pseudogene | 1,68 | 0,75 | 8,46E-05 |
| ***Slc1a4*** | solute carrier family 1 (glutamate/neutral amino acid transporter), member 4 | 1,67 | 0,74 | 4,46E-14 |
| ***Selenbp1*** | selenium binding protein 1 | 1,66 | 0,73 | 3,55E-03 |
| ***P4ha2*** | procollagen-proline, 2-oxoglutarate 4-dioxygenase (proline 4-hydroxylase), alpha II polypeptide | 1,66 | 0,73 | 8,29E-31 |
| ***2500002B13Rik*** | RIKEN cDNA 2500002B13 gene | 1,64 | 0,72 | 3,41E-03 |
| ***Egln3*** | egl-9 family hypoxia-inducible factor 3 | 1,63 | 0,71 | 9,31E-03 |
| ***Dusp10*** | dual specificity phosphatase 10 | 1,63 | 0,70 | 5,02E-03 |
| ***Nfil3*** | nuclear factor, interleukin 3, regulated | 1,63 | 0,70 | 7,21E-08 |
| ***Slc16a3*** | solute carrier family 16 (monocarboxylic acid transporters), member 3 | 1,60 | 0,68 | 1,54E-06 |
| ***Gys1*** | glycogen synthase 1, muscle | 1,60 | 0,68 | 4,40E-06 |
| ***Pkp2*** | plakophilin 2 | 1,59 | 0,67 | 2,94E-05 |
| ***Sesn2*** | sestrin 2 | 1,58 | 0,66 | 1,03E-22 |
| ***Hoxb9*** | homeobox B9 | 1,58 | 0,66 | 1,86E-10 |
| ***Btg2*** | BTG anti-proliferation factor 2 | 1,58 | 0,66 | 2,40E-04 |
| ***Sap30*** | sin3 associated polypeptide | 1,57 | 0,65 | 7,73E-10 |
| ***Pdk1*** | pyruvate dehydrogenase kinase, isoenzyme 1 | 1,57 | 0,65 | 3,09E-03 |
| ***Rnf19a*** | ring finger protein 19A | 1,55 | 0,64 | 3,17E-06 |
| ***Rora*** | RAR-related orphan receptor alpha | 1,55 | 0,63 | 1,32E-07 |
| ***Rhob*** | ras homolog family member B | 1,55 | 0,63 | 7,80E-15 |
| ***Ppp1r15a*** | protein phosphatase 1, regulatory subunit 15A | 1,54 | 0,63 | 1,34E-11 |
| ***Slc2a1*** | solute carrier family 2 (facilitated glucose transporter), member 1 | 1,53 | 0,62 | 1,60E-07 |
| ***Plk3*** | polo like kinase 3 | 1,53 | 0,61 | 3,40E-06 |
| ***Arid5a*** | AT-rich interaction domain 5A | 1,51 | 0,60 | 5,52E-09 |
| ***Egln1*** | egl-9 family hypoxia-inducible factor 1 | 1,51 | 0,60 | 1,17E-06 |
| ***Jund*** | jun D proto-oncogene | 1,51 | 0,59 | 2,29E-05 |
| ***Nfkbiz*** | nuclear factor of kappa light polypeptide gene enhancer in B cells inhibitor, zeta | 1,50 | 0,58 | 1,17E-04 |
| ***Kif21b*** | kinesin family member 21B | 1,49 | 0,58 | 3,17E-25 |
| ***Prkab2*** | protein kinase, AMP-activated, beta 2 non-catalytic subunit | 1,47 | 0,56 | 1,61E-06 |
| ***Rcor2*** | REST corepressor 2 | 1,47 | 0,55 | 1,28E-07 |
| ***Tbl2*** | transducin (beta)-like 2 | 1,45 | 0,54 | 1,90E-16 |

| Gene | Entrez Gene Name | FC | Log$_2$FC | $p_{adj}$ value |
|---|---|---|---|---|
| ***Fam102a*** | estrogen-induced osteoclastogenesis regulator 1 | 1,45 | 0,54 | 9,01E-09 |
| ***Zfp395*** | zinc finger protein 395 | 1,45 | 0,54 | 6,65E-07 |
| ***Dusp16*** | dual specificity phosphatase 16 | 1,45 | 0,54 | 1,05E-12 |
| ***Hmox1*** | heme oxygenase 1 | 1,45 | 0,53 | 1,25E-03 |
| ***Mxi1*** | MAX interactor 1, dimerization protein | 1,44 | 0,53 | 8,38E-09 |
| ***Hyal1*** | hyaluronoglucosaminidase 1 | 1,44 | 0,53 | 1,73E-07 |
| ***Csrnp1*** | cysteine-serine-rich nuclear protein 1 | 1,44 | 0,52 | 3,40E-05 |
| ***Phldb3*** | pleckstrin homology like domain, family B, member 3 | 1,43 | 0,52 | 3,22E-02 |
| ***Slc7a11*** | solute carrier family 7 (cationic amino acid transporter, y+ system), member 11 | 1,43 | 0,51 | 3,40E-06 |
| ***9330188P03Rik*** | RIKEN cDNA 9330188P03 gene | 1,42 | 0,50 | 6,18E-05 |
| ***Kdm5b*** | lysine demethylase 5B | 1,41 | 0,50 | 6,49E-06 |
| ***Dubr*** | Dppa2 upstream binding RNA | 1,41 | 0,49 | 1,51E-03 |
| ***Siah2*** | siah E3 ubiquitin protein ligase 2 | 1,40 | 0,49 | 7,11E-05 |
| ***Zfp36*** | zinc finger protein 36 | 1,39 | 0,47 | 8,32E-04 |
| ***Higd1a*** | HIG1 domain family, member 1A | 1,39 | 0,47 | 1,54E-03 |
| ***Ankzf1*** | ankyrin repeat and zinc finger domain containing 1 | 1,38 | 0,47 | 2,72E-06 |
| ***Plod2*** | procollagen lysine, 2-oxoglutarate 5-dioxygenase 2 | 1,38 | 0,47 | 1,07E-03 |
| ***Khnyn*** | KH and NYN domain containing | 1,38 | 0,47 | 6,49E-06 |
| ***Vhl*** | von Hippel-Lindau tumor suppressor | 1,38 | 0,47 | 2,91E-03 |
| ***Prelid2*** | PRELI domain containing 2 | 1,38 | 0,47 | 2,63E-03 |
| ***Sertad1*** | SERTA domain containing 1 | 1,38 | 0,47 | 5,41E-04 |
| ***Plaur*** | plasminogen activator, urokinase receptor | 1,38 | 0,46 | 2,08E-05 |
| ***Filip1l*** | filamin A interacting protein 1-like | 1,37 | 0,46 | 1,49E-05 |
| ***Gm9790*** | predicted gene 9790 | 1,37 | 0,46 | 7,11E-05 |
| ***Rassf1*** | Ras association (RalGDS/AF-6) domain family member 1 | 1,36 | 0,45 | 1,20E-03 |
| ***Pnrc1*** | proline-rich nuclear receptor coactivator 1 | 1,36 | 0,44 | 1,85E-06 |
| ***Per1*** | period circadian clock 1 | 1,36 | 0,44 | 1,35E-07 |
| ***Soat1*** | sterol O-acyltransferase 1 | 1,35 | 0,43 | 9,75E-04 |
| ***P4ha1*** | procollagen-proline, 2-oxoglutarate 4-dioxygenase (proline 4-hydroxylase), alpha II polypeptide | 1,35 | 0,43 | 6,52E-12 |
| ***Adamts1*** | ADAM metallopeptidase with thrombospondin type 1 motif 5 | 1,35 | 0,43 | 5,64E-20 |
| ***Eno2*** | enolase 2, gamma neuronal | 1,34 | 0,43 | 4,78E-05 |
| ***Pfkp*** | phosphofructokinase, platelet | 1,34 | 0,42 | 2,25E-03 |
| ***Kdelr3*** | KDEL (Lys-Asp-Glu-Leu) endoplasmic reticulum protein retention receptor 3 | 1,33 | 0,41 | 1,70E-04 |
| ***Irak2*** | interleukin-1 receptor-associated kinase 2 | 1,33 | 0,41 | 1,03E-02 |
| ***Apold1*** | apolipoprotein L domain containing 1 | 1,33 | 0,41 | 1,28E-03 |
| ***Kdm3a*** | lysine (K)-specific demethylase 3A | 1,32 | 0,40 | 5,43E-05 |
| ***Noct*** | nocturnin | 1,32 | 0,40 | 8,46E-05 |
| ***Rusc2*** | RUN and SH3 domain containing 2 | 1,32 | 0,40 | 1,94E-12 |

| Gene | Entrez Gene Name | FC | $Log_2FC$ | $p_{adj}$ value |
|---|---|---|---|---|
| ***Dusp2*** | dual specificity phosphatase 2 | 1,32 | 0,40 | 1,20E-03 |
| ***Jmjd6*** | jumonji domain containing 6 | 1,31 | 0,39 | 2,07E-11 |
| ***Otud1*** | OTU domain containing 1 | 1,31 | 0,39 | 2,40E-04 |
| ***Naa80*** | N(alpha)-acetyltransferase 80, NatH catalytic subunit | 1,31 | 0,39 | 5,78E-07 |
| ***Sema6b*** | sema domain, transmembrane domain (TM), and cytoplasmic domain, (semaphorin) 6B | 1,31 | 0,39 | 4,28E-05 |
| ***Foxo3*** | forkhead box O3 | 1,31 | 0,39 | 2,92E-08 |
| ***Kdm4b*** | lysine (K)-specific demethylase 4B | 1,31 | 0,39 | 1,77E-07 |
| ***Skil*** | SKI-like | 1,30 | 0,38 | 1,28E-03 |
| ***Grhpr*** | glyoxylate reductase/hydroxypyruvate reductase | 1,30 | 0,38 | 3,33E-06 |
| ***Ubald1*** | UBA-like domain containing 1 | 1,30 | 0,38 | 2,62E-04 |
| ***Pck2*** | phosphoenolpyruvate carboxykinase 2 (mitochondrial) | 1,30 | 0,38 | 2,71E-05 |
| ***Gpt2*** | glutamic pyruvate transaminase (alanine aminotransferase) 2 | 1,30 | 0,38 | 6,59E-05 |
| ***Inafm2*** | InaF motif containing 2 | 1,30 | 0,37 | 2,83E-04 |
| ***Bmt2*** | S-adenosylmethionine sensor upstream of mTORC1 | 1,30 | 0,37 | 3,89E-08 |
| ***Pvr*** | nectin cell adhesion molecule 2 | 1,29 | 0,37 | 7,09E-07 |
| ***Gm49759*** | predicted gene, 49759 | 1,29 | 0,37 | 7,88E-11 |
| ***Pfkl*** | phosphofructokinase, liver, B-type | 1,29 | 0,37 | 1,05E-02 |
| ***Maml1*** | mastermind like transcriptional coactivator 1 | 1,29 | 0,37 | 7,68E-07 |
| ***Map3k1*** | mitogen-activated protein kinase kinase kinase 1 | 1,29 | 0,36 | 1,23E-05 |
| ***Atf4*** | activating transcription factor 4 | 1,28 | 0,36 | 1,82E-18 |
| ***Arl14ep*** | ADP-ribosylation factor-like 14 effector protein | 1,28 | 0,36 | 6,30E-05 |
| ***Srf*** | serum response factor | 1,27 | 0,35 | 7,11E-05 |
| ***Tnfaip2*** | tumor necrosis factor, alpha-induced protein 2 | 1,27 | 0,34 | 1,06E-06 |
| ***Zswim4*** | zinc finger SWIM-type containing 4 | 1,27 | 0,34 | 2,00E-02 |
| ***Kdm6b*** | KDM1 lysine (K)-specific demethylase 6B | 1,27 | 0,34 | 8,47E-06 |
| ***Junb*** | jun B proto-oncogene | 1,26 | 0,34 | 4,08E-02 |
| ***Kctd13*** | potassium channel tetramerisation domain containing 13 | 1,26 | 0,33 | 4,29E-02 |
| ***Bnip3l*** | BCL2/adenovirus E1B interacting protein 3-like | 1,26 | 0,33 | 1,93E-06 |
| ***Stk40*** | serine/threonine kinase 40 | 1,26 | 0,33 | 1,78E-04 |
| ***Ppme1*** | protein phosphatase methylesterase 1 | 1,26 | 0,33 | 1,76E-06 |
| ***Errfi1*** | ERBB receptor feedback inhibitor 1 | 1,26 | 0,33 | 4,52E-03 |
| ***Cebpg*** | CCAAT/enhancer binding protein gamma | 1,26 | 0,33 | 1,27E-07 |
| ***Mthfd2*** | methylenetetrahydrofolate dehydrogenase (NAD+ dependent), methenyltetrahydrofolate cyclohydrolase | 1,26 | 0,33 | 3,02E-11 |
| ***Pde4dip*** | phosphodiesterase 4D interacting protein (myomegalin) | 1,26 | 0,33 | 4,86E-04 |
| ***Slc37a4*** | solute carrier family 37 (glucose-6-phosphate transporter), member 4 | 1,25 | 0,33 | 2,73E-03 |
| ***Clcn3*** | chloride channel, voltage-sensitive 3 | 1,25 | 0,33 | 1,06E-08 |
| ***Unc5b*** | unc-5 netrin receptor B | 1,25 | 0,32 | 1,24E-02 |

| Gene | Entrez Gene Name | FC | Log$_2$FC | $p_{adj}$ value |
|---|---|---|---|---|
| ***Bnip3l-ps*** | BCL2/adenovirus E1B interacting protein 3-like, pseudogene | 1,25 | 0,32 | 3,97E-02 |
| ***Gzf1*** | GDNF-inducible zinc finger protein 1 | 1,25 | 0,32 | 4,69E-06 |
| ***Tet1*** | tet methylcytosine dioxygenase 1 | 1,24 | 0,31 | 2,69E-02 |
| ***Pgm1*** | phosphoglucomutase 2 | 1,24 | 0,31 | 4,60E-02 |
| ***Nr4a3*** | nuclear receptor subfamily 4, group A, member 3 | 1,24 | 0,31 | 9,07E-05 |
| ***Herpud1*** | homocysteine-inducible, endoplasmic reticulum stress-inducible, ubiquitin-like domain member 1 | 1,24 | 0,30 | 1,10E-09 |
| ***Ldlrap1*** | low density lipoprotein receptor adaptor protein 1 | 1,23 | 0,30 | 4,29E-03 |
| ***Eif4ebp1*** | eukaryotic translation initiation factor 4E binding protein 1 | 1,23 | 0,30 | 2,75E-05 |
| ***Vgll4*** | vestigial like family member 4 | 1,23 | 0,30 | 1,89E-10 |
| ***Lonrf1*** | LON peptidase N-terminal domain and ring finger 1 | 1,23 | 0,30 | 1,33E-02 |
| ***Fbxo10*** | F-box protein 10 | 1,23 | 0,30 | 1,78E-03 |
| ***Gla*** | galactosidase, alpha | 1,23 | 0,29 | 1,67E-02 |
| ***Ugdh*** | UDP-glucose dehydrogenase | 1,22 | 0,29 | 1,11E-04 |
| ***Slc25a25*** | solute carrier family 25 (mitochondrial carrier, phosphate carrier), member 25 | 1,22 | 0,29 | 1,47E-02 |
| ***Spry2*** | sprouty RTK signaling antagonist 2 | 1,22 | 0,29 | 3,60E-02 |
| ***Klf6*** | Kruppel-like transcription factor 6 | 1,22 | 0,29 | 1,56E-05 |
| ***Fam219a*** | family with sequence similarity 219, member A | 1,22 | 0,28 | 1,23E-03 |
| ***Itpk1*** | inositol 1,3,4-triphosphate 5/6 kinase | 1,22 | 0,28 | 7,70E-04 |
| ***Ppargc1a*** | peroxisome proliferative activated receptor, gamma, coactivator 1 alpha | 1,21 | 0,28 | 1,43E-04 |
| ***Cttnbp2nl*** | CTTNBP2 N-terminal like | 1,21 | 0,28 | 1,06E-03 |
| ***Nampt*** | nicotinamide phosphoribosyltransferase | 1,21 | 0,28 | 2,91E-02 |
| ***Cars*** | cysteinyl-tRNA synthetase 1 | 1,21 | 0,28 | 7,47E-05 |
| ***Tlcd3a*** | TLC domain containing 3A | 1,21 | 0,28 | 5,76E-03 |
| ***Acvrl1*** | activin A receptor, type II-like 1 | 1,21 | 0,28 | 4,49E-06 |
| ***Cnn2*** | calponin 2 | 1,21 | 0,28 | 7,05E-03 |
| ***Gm12715*** | actin, gamma, cytoplasmic 1 pseudogene | 1,21 | 0,27 | 6,63E-05 |
| ***Sdc4*** | syndecan 4 | 1,21 | 0,27 | 2,62E-04 |
| ***Rnf217*** | ring finger protein 217 | 1,21 | 0,27 | 1,26E-02 |
| ***Rsbn1*** | rosbin, round spermatid basic protein 1 | 1,21 | 0,27 | 4,86E-02 |
| ***Phlda1*** | pleckstrin homology like domain, family A, member 1 | 1,21 | 0,27 | 3,20E-06 |
| ***St3gal4*** | ST3 beta-galactoside alpha-2,3-sialyltransferase 4 | 1,20 | 0,27 | 7,09E-07 |
| ***Kdm7a*** | lysine (K)-specific demethylase 7A | 1,20 | 0,27 | 2,40E-02 |
| ***Gtpbp2*** | GTP binding protein 2 | 1,20 | 0,26 | 1,63E-04 |
| ***Alkbh5*** | alkB homolog 5, RNA demethylase | 1,20 | 0,26 | 7,09E-07 |
| ***Frmd3*** | FERM domain containing 3 | 1,20 | 0,26 | 3,82E-02 |
| ***Samd4b*** | sterile alpha motif domain containing 4B | 1,20 | 0,26 | 4,83E-04 |
| ***Ier2*** | immediate early response 2 | 1,20 | 0,26 | 1,46E-02 |

| Gene | Entrez Gene Name | FC | $Log_2FC$ | $p_{adj}$ value |
|---|---|---|---|---|
| ***Zbtb21*** | zinc finger and BTB domain containing 21 | 1,20 | 0,26 | 1,59E-02 |
| ***Appl2*** | adaptor protein, phosphotyrosine interaction, PH domain and leucine zipper containing 2 | 1,20 | 0,26 | 1,44E-05 |
| ***Adamts4*** | ADAM metallopeptidase with thrombospondin type 1 motif 4 | 1,19 | 0,25 | 6,95E-04 |
| ***Rlf*** | ral guanine nucleotide dissociation stimulator-like 2 | 1,19 | 0,25 | 1,81E-05 |
| ***Gm8399*** | actin, gamma, cytoplasmic 1 pseudogene | 1,19 | 0,25 | 2,82E-02 |
| ***Gab2*** | growth factor receptor bound protein 2-associated protein 2 | 1,19 | 0,25 | 4,05E-03 |
| ***Ccdc115*** | coiled-coil domain containing 115 | 1,19 | 0,25 | 6,38E-03 |
| ***Zfp292*** | zinc finger protein 292 | 1,18 | 0,24 | 9,31E-03 |
| ***Bcar1*** | breast cancer anti-estrogen resistance 1 | 1,18 | 0,24 | 1,41E-03 |
| ***Cdkn1a*** | cyclin dependent kinase inhibitor 1A | 1,18 | 0,24 | 6,14E-07 |
| ***Peli1*** | pellino 1 | 1,18 | 0,24 | 2,96E-02 |
| ***Aldh18a1*** | aldehyde dehydrogenase 18 family, member A1 | 1,18 | 0,24 | 2,45E-03 |
| ***Trmt10c*** | tRNA methyltransferase 10C, mitochondrial RNase P subunit | 1,18 | 0,24 | 1,30E-02 |
| ***Dusp4*** | dual specificity phosphatase 4 | 1,18 | 0,24 | 1,04E-06 |
| ***Zfp654*** | zinc finger protein 654 | 1,18 | 0,24 | 1,66E-03 |
| ***Mafk*** | v-maf musculoaponeurotic fibrosarcoma oncogene family, protein K (avian) | 1,18 | 0,24 | 5,02E-03 |
| ***Gpi1*** | phosphatidylinositol glycan anchor biosynthesis, class Q | 1,18 | 0,24 | 4,61E-02 |
| ***Sars*** | seryl-tRNA synthetase 1 | 1,18 | 0,23 | 9,52E-07 |
| ***Ip6k2*** | inositol hexaphosphate kinase 2 | 1,17 | 0,23 | 3,51E-02 |
| ***Espn*** | espin | 1,17 | 0,23 | 3,46E-02 |
| ***Tiparp*** | TCDD-inducible poly(ADP-ribose) polymerase | 1,17 | 0,23 | 1,46E-02 |
| ***Arhgef2*** | Rho/Rac guanine nucleotide exchange factor 2 | 1,17 | 0,23 | 1,91E-07 |
| ***Actg1*** | actin, gamma, cytoplasmic 1 | 1,17 | 0,23 | 1,51E-03 |
| ***Asns*** | asparagine synthetase | 1,17 | 0,23 | 1,28E-03 |
| ***Arrdc3*** | arrestin domain containing 3 | 1,17 | 0,22 | 2,19E-02 |
| ***Mef2d*** | myocyte enhancer factor 2D | 1,17 | 0,22 | 1,80E-05 |
| ***Rab31*** | RAB31, member RAS oncogene family | 1,17 | 0,22 | 1,06E-02 |
| ***Gpi-ps*** | glucose-6-phosphate isomerase, pseudogene | 1,16 | 0,22 | 7,47E-03 |
| ***Cdc42se1*** | CDC42 small effector 1 | 1,16 | 0,22 | 6,00E-04 |
| ***Irf2bp2*** | interferon regulatory factor 2 binding protein 2 | 1,16 | 0,21 | 4,69E-02 |
| ***Usp22*** | ubiquitin specific peptidase 22 | 1,16 | 0,21 | 3,13E-03 |
| ***Arih2*** | ariadne RBR E3 ubiquitin protein ligase 2 | 1,16 | 0,21 | 2,63E-03 |
| ***2310022B05Rik*** | RIKEN cDNA 2310022B05 gene | 1,15 | 0,21 | 5,84E-03 |
| ***Dnajc5*** | DnaJ heat shock protein family (Hsp40) member C5 | 1,15 | 0,21 | 2,40E-04 |
| ***Spty2d1*** | SPT2 chromatin protein domain containing 1 | 1,15 | 0,21 | 3,53E-02 |
| ***Neat1*** | nuclear paraspeckle assembly transcript 1 (non-protein coding) | 1,15 | 0,21 | 8,14E-06 |
| ***Gtf2e2*** | general transcription factor II E, polypeptide 2 (beta subunit) | 1,15 | 0,20 | 1,22E-02 |

| Gene | Entrez Gene Name | FC | $Log_2FC$ | $p_{adj}$ value |
|---|---|---|---|---|
| ***Vasp*** | vasodilator-stimulated phosphoprotein | 1,15 | 0,20 | 1,14E-04 |
| ***Zfp655*** | zinc finger protein 655 | 1,15 | 0,20 | 2,13E-03 |
| ***Ube2o*** | ubiquitin-conjugating enzyme E2O | 1,15 | 0,20 | 3,13E-03 |
| ***Cdkn1b*** | cyclin dependent kinase inhibitor 1B | 1,14 | 0,19 | 5,24E-03 |
| ***Wsb1*** | WD repeat and SOCS box-containing 1 | 1,14 | 0,19 | 1,10E-03 |
| ***Kmt2e*** | lysine (K)-specific methyltransferase 2E | 1,14 | 0,19 | 5,52E-04 |
| ***Ufsp2*** | UFM1-specific peptidase 2 | 1,14 | 0,19 | 4,59E-02 |
| ***Usp36*** | ubiquitin specific peptidase 36 | 1,14 | 0,19 | 1,22E-02 |
| ***Cmtm3*** | CKLF-like MARVEL transmembrane domain containing 3 | 1,14 | 0,19 | 9,94E-03 |
| ***Ugp2*** | UDP-glucose pyrophosphorylase 2 | 1,14 | 0,19 | 3,61E-03 |
| ***Cflar*** | CASP8 and FADD-like apoptosis regulator | 1,13 | 0,18 | 3,55E-03 |
| ***Rhou*** | ras homolog family member U | 1,13 | 0,18 | 4,82E-02 |
| ***Snx33*** | sorting nexin 33 | 1,13 | 0,18 | 2,12E-02 |
| ***Cep170*** | centrosomal protein 170 | 1,13 | 0,18 | 2,73E-03 |
| ***Pgam1*** | phosphoglycerate mutase 1 | 1,13 | 0,18 | 3,20E-02 |
| ***Abl2*** | ABL proto-oncogene 2, non-receptor tyrosine kinase | 1,13 | 0,18 | 1,46E-02 |
| ***Sh3pxd2b*** | SH3 and PX domains 2B | 1,13 | 0,17 | 1,67E-02 |
| ***Gopc*** | golgi associated PDZ and coiled-coil motif containing | 1,13 | 0,17 | 2,57E-02 |
| ***H1f0*** | H1.0 linker histone | 1,13 | 0,17 | 6,35E-03 |
| ***Strip1*** | striatin interacting protein 1 | 1,13 | 0,17 | 2,41E-02 |
| ***G2e3*** | G2/M-phase specific E3 ubiquitin ligase | 1,13 | 0,17 | 4,29E-02 |
| ***Creld1*** | cysteine-rich with EGF-like domains 1 | 1,12 | 0,17 | 3,50E-02 |
| ***Kdm1a*** | lysine (K)-specific demethylase 1A | 1,12 | 0,17 | 2,35E-02 |
| ***Map2k1*** | mitogen-activated protein kinase kinase 1 | 1,12 | 0,17 | 1,26E-03 |
| ***Rcor1*** | REST corepressor 2 | 1,12 | 0,17 | 3,02E-03 |
| ***Gas2l3*** | growth arrest-specific 2 like 3 | 1,12 | 0,17 | 6,23E-03 |
| ***Prdm16*** | PR domain containing 16 | 1,12 | 0,17 | 1,37E-03 |
| ***Mef2a*** | myocyte enhancer factor 2A | 1,12 | 0,16 | 3,53E-02 |
| ***Ifrd1*** | interferon-related developmental regulator 1 | 1,12 | 0,16 | 2,27E-02 |
| ***Shc4*** | SHC (Src homology 2 domain containing) family, member 4 | 1,12 | 0,16 | 2,63E-02 |
| ***Slc38a2*** | solute carrier family 38, member 2 | 1,12 | 0,16 | 2,49E-04 |
| ***Lonp1*** | lon peptidase 1, mitochondrial | 1,12 | 0,16 | 1,46E-02 |
| ***Slc7a1*** | solute carrier family 7 (cationic amino acid transporter, y+ system), member 3 | 1,11 | 0,16 | 1,13E-04 |
| ***Acox1*** | acyl-Coenzyme A oxidase 1, palmitoyl | 1,11 | 0,15 | 3,51E-02 |
| ***Nars*** | asparaginyl-tRNA synthetase 1 | 1,11 | 0,15 | 1,07E-03 |
| ***Mrs2*** | MRS2 magnesium transporter | 1,11 | 0,15 | 1,26E-02 |
| ***Ccdc137*** | coiled-coil domain containing 137 | 1,11 | 0,15 | 2,53E-02 |
| ***Aars*** | alanyl-tRNA synthetase 1 | 1,11 | 0,15 | 1,07E-03 |
| ***Mia2*** | MIA SH3 domain ER export factor 2 | 1,11 | 0,15 | 4,05E-02 |

| Gene | Entrez Gene Name | FC | $Log_2FC$ | $p_{adj}$ value |
|---|---|---|---|---|
| ***Ldha*** | lactate dehydrogenase A | 1,11 | 0,15 | 1,60E-02 |
| ***Slc3a2*** | solute carrier family 3 (activators of dibasic and neutral amino acid transport), member 2 | 1,11 | 0,15 | 1,22E-03 |
| ***Rnf181*** | ring finger protein 181 | 1,11 | 0,14 | 1,24E-02 |
| ***Smcr8*** | Smith-Magenis syndrome chromosome region, candidate 8 homolog (human) | 1,10 | 0,14 | 3,51E-02 |
| ***Nckap5l*** | NCK-associated protein 5-like | 1,10 | 0,14 | 2,69E-02 |
| ***Slc7a5*** | solute carrier family 7 (cationic amino acid transporter, y+ system), member 5 | 1,10 | 0,14 | 2,40E-03 |
| ***Fyttd1*** | forty-two-three domain containing 1 | 1,10 | 0,14 | 3,28E-02 |
| ***Ndrg1*** | N-myc downstream regulated gene 1 | 1,10 | 0,14 | 4,50E-02 |
| ***Szrd1*** | SUZ RNA binding domain containing 1 | 1,10 | 0,14 | 1,23E-03 |
| ***Tpm4*** | tropomyosin 4 | 1,10 | 0,14 | 3,73E-02 |
| ***Stat6*** | signal transducer and activator of transcription 6 | 1,09 | 0,13 | 2,52E-02 |
| ***Pax3*** | paired box 3 | 1,09 | 0,13 | 4,50E-02 |
| ***Litaf*** | LPS-induced TN factor | 1,09 | 0,12 | 3,86E-02 |
| ***Mknk2*** | MAP kinase-interacting serine/threonine kinase 2 | 1,09 | 0,12 | 1,26E-02 |
| ***Ghitm*** | growth hormone inducible transmembrane protein | 1,08 | 0,11 | 2,96E-02 |

**Supplementary Table 3. List of primers used in RT-qPCR analysis.**

| Gene | Forward (5’ → 3’) | Reverse (3’ → 5’) |
|---|---|---|
| ***Fos*** | ACTTCGACCATGATGTTCTCG | GCTGGGGAATGGTAGTAGGAA |
| ***Jun*** | GGGTGCCAACTCATGCTAAC | CGCAACCAGTCAAGTTCTCA |
| ***Jund*** | GACCCTCAAAAGCCAGAACA | GACGTGGCTGAGGACTTTCT |
| ***Atf3*** | AACAGAGGATGGACGACACC | GCCTTCATTGTGTGACGTTG |
| ***Hif1a*** | CCCAATGGATGATGATTTCC | GGTCTGCTGGAACCCAGTAA |
| ***P38a*** | GCTGAACAAAGGGAGAGACGA | GGGCCTTGATGACTTGGTTTG |
| ***Slc2a1*** | AAACTCCAGTCAGCATTTTCATAGG | CCACGGAGTTGTAAAATATGCTTGT |
| ***Hk2*** | ATGTACTTGGGCGAGATTGTG | GTTTCGAAGATTCCCCTTGTC |
| ***Eno2*** | GGGCACTCTACCAGGACTTT | TCAGGTCATCGCCCACTATC |
| ***Rpl19*** | GACCAAGGAAGCACGAAAGC | CAGGCCGCTATGTACAGACA |
